# An illustration of reproducibility in neuroscience research in the absence of selective reporting

**DOI:** 10.1101/866301

**Authors:** Xiang-Zhen Kong, ENIGMA Laterality Working Group, Clyde Francks

**Affiliations:** Language and Genetics Department, Max Planck Institute for Psycholinguistics, Nijmegen, The Netherlands; Donders Institute for Brain, Cognition and Behaviour, Radboud University, Nijmegen, The Netherlands

**Keywords:** Reproducibility, Publication bias, *P*-hacking, Team science, Multi-site collaboration

## Abstract

The problem of poor reproducibility of scientific findings has received much attention over recent years, in a variety of fields including psychology and neuroscience. The problem has been partly attributed to publication bias and unwanted practices such as *p*-hacking. Low statistical power in individual studies is also understood to be an important factor. In a recent multi-site collaborative study, we mapped brain anatomical left-right asymmetries for regional measures of surface area and cortical thickness, in 99 MRI datasets from around the world, for a total of over 17,000 participants. In the present study, we re-visited these hemispheric effects from the perspective of reproducibility. Within each dataset, we considered that an effect had been reproduced when it matched the meta-analytic effect from the 98 other datasets, in terms of effect direction and uncorrected significance at *p*<0.05. In this sense, the results within each dataset were viewed as coming from separate studies in an ‘ideal publishing environment’, i.e. free from selective reporting and *p* hacking. We found an average reproducibility rate per dataset, over all effects, of 63.2% (*SD* = 22.9%, min = 22.2%, max = 97.0%). As expected, reproducibility was higher for larger effects and in larger datasets. There is clearly substantial room to improve reproducibility in brain MRI research through increasing statistical power. These findings constitute an empirical illustration of reproducibility in the absence of publication bias or *p* hacking, when assessing realistic biological effects in heterogeneous neuroscience data, and given typically-used sample sizes.

---

The issue of reproducibility has received considerable attention in a variety of fields including medicine (Prinz et al. 2011), psychology (Aarts et al. 2015; Klein et al. 2014) and neuroscience (Button et al. 2013). Poor reproducibility has been partly attributed to reporting bias and problematic practices such as selective reporting of outcomes (i.e. *p*-hacking) (Aarts et al. 2015; Baker 2016; Bakker et al. 2012; Ioannidis et al. 2014; Ioannidis 2005; Ioannidis 2008; John et al. 2012; Simmons et al. 2011). This situation has resulted in multiple calls for more reproducible research (e.g., (Benjamin et al. 2018; Button et al. 2013; Klein et al. 2018a; Poldrack et al. 2017; Valentin Amrhein 2017)). For example, the Open Science Framework has been set up as a free and open source project management resource for researchers across the entire study cycle. In addition, the Transparency and Openness Promotion (TOP) Guidelines (Nosek et al. 2015) have been proposed to improve the quality and credibility of scientific literature. In neuroimaging studies, problems such as flexibility in data analysis have been widely discussed, and best practices have been proposed to ensure that neuroimaging studies can produce meaningful and reliable results (Poldrack et al. 2017). The reproducibility rate was not found to correlate with levels of experience and expertise of study authors, in a replication study of previous findings in psychology (Aarts et al. 2015), which suggests that some practices will not improve merely through training, and that other factors influence reproducibility within the current research convention.

Among these factors, low statistical power is now well understood to contribute to the reproducibility problem (Button et al. 2013; Ioannidis 2005). The positive predictive value (PPV), i.e. the probability that a ‘positive’ research finding reflects a true effect, has been formulated as a function of the prior probability of the effect being real (*R*, the pre-study odds), the statistical power of the study (*1*−*β; β* is the type II error), and the level of statistical significance required (*α*; *α* is the type I error, e.g., 0.05 or 0.01): *PPV = (1 − β)R/((1− β)R+α))* (Button et al. 2013; Ioannidis 2005). For example, it is evident that a research finding is more likely true than false (i.e., *PPV* > 50%) if *(1− β)R* > *α*. However, in many cases the true effect size is unknown *a priori*, and/or the pre-study odds are unknown. This problem is then further complicated by potentially selective reporting.

The present study aimed to demonstrate reproducibility in a real-world setting, where *a priori* knowledge of the statistical power and pre-study odds was not necessary, and in the absence of selective reporting. We also aimed to show how reproducibility changes with the real effect size and sample size used, which together determine the statistical power. If we wanted to address this question with actual papers in the literature, an ideal publishing environment, free from selective reporting, would first need to be established. This seems impossible in the current era, because many journals and scientists are incentivized to report statistically significant results, while leaving non-significant findings unpublished (known as the *file draw* effect). Here we leveraged summary statistics from a study performed via a worldwide collaborative network, known as the Enhancing NeuroImaging Genetics through Meta-Analysis (ENIGMA) consortium (Thompson et al. 2014a), to estimate reproducibility when there is no reporting bias. Briefly, the ENIGMA consortium allows researchers worldwide to work together on the same questions in neuroscience, genetics, and psychiatry. Analysis plans and scripts are prepared by a central site before running any analysis, and then sent out to each separate site to run on their own dataset. Finally, outputs from every dataset are sent back to the central site and synthesized by the application of meta-analysis methodology. Thus, we can consider outputs from each dataset as being from an “ideal reporting environment”, free from reporting bias, or other potentially problematic practices such as *p*-hacking. If we assume the overall meta-effect size to represent the ‘true’ effect size, in this way we have access to a real-world setting for examining reproducibility in the absence of selective reporting, which can provide a useful illustration of how consistently some realistic effects can be detected when surveying cohorts worldwide.

Here, we made use of summary statistics from the ENIGMA project on mapping cerebral cortical left-right asymmetry (Kong et al. 2018), which involved 99 datasets and a total of 17,141 participants worldwide (see Materials and Methods). We focused our analyses on hemispheric effects on cerebral cortical thickness and surface area measures. The human brain is subtly asymmetrical on the left-right axis, so that homologous regions in the two hemispheres can differ with respect to their cortical thickness and surface area (Kong et al. 2018). Specifically, we analyzed thickness and area measures for each of 34 brain regions based on the Desikan-Killany atlas from *FreeSurfer*, as well as entire hemisphere-level average thickness and total area (Fischl 2012), for a total of 70 left-right hemispheric effects. To simulate an ideal publishing environment, here we considered each asymmetry as one single research question (e.g., does the parahippocampal gyrus show left-right asymmetrical thickness, on average, in the human brain?). In other words, in the context of the present study of reproducibility, we effectively ran 70 research questions about asymmetry, each repeated 99 times in different datasets, but with different sample sizes and scanning equipment and parameters (although image processing with *Freesurfer* was harmonized across sites). These data allowed us to assess reproducibility in an “ideal publishing environment”, but also in a real-world context of dataset heterogeneity. We also examined how reproducibility changed in relation to effect sizes, and the sample sizes of individual datasets. Although our empirical illustration focused on the field of human brain MRI research, the implications are broadly applicable to many fields.

## Materials and Methods

### Datasets

In this study, we used the publicly available statistical outputs of the ENIGMA cortical asymmetry project (http://conxz.net/neurohemi/) (Kong et al. 2018). These datasets comprised 17,141 healthy participants in total from 99 datasets, each of which showed different sex, age and handedness distributions, and were from diverse ethnic backgrounds. Participants were drawn from the general population or were healthy controls from clinical studies. **Table S1** gives summary information for each dataset. All local institutional review boards permitted the use of extracted measures from the anonymized data.

Here, we focused the reproducibility analyses on the hemispheric effects on paired left-right measures of cortical thickness and surface area, for 34 brain regions based on the Desikan-Killany atlas from *FreeSurfer* (Fischl 2012), as well as entire hemisphere-level average thickness and total surface area. See Kong et al., 2018 for details about the neuroimaging processing and quality control. Briefly, images were acquired using scanners of different field strengths (1.5T and 3T) and all images were analyzed using the automated and validate pipeline “recon-all” implemented in *FreeSurfer* (Fischl 2012), although different software versions were possible (version 5.0, 5.1, and 5.3) (Table S1). Together with the varying demographic composition as summarized above, these datasets illustrate the heterogeneity that is a feature of real-world neuroimaging data.

For each dataset and paired left-right measure, paired t-tests were used to assess inter-hemispheric differences, and Cohen’s *d* was calculated based on each paired t-test, to estimate the effect size. In the procedure, analysis plans and scripts were prepared by a central site and sent out to each dataset’s own site for running the analysis, and finally all outputs for every dataset were sent back to the central site.

### Estimation of the ‘True’ Effects

Given a lack of consistency of previous brain asymmetry findings in the earlier literature, Kong et al. 2018 performed the largest ever study of this issue, based on at least an order of magnitude more participants than any previous study (Kong et al. 2018). For each hemispheric effect on regional or total thickness or surface area, the outputs from each separate dataset were combined using inverse variance-weighted random-effect meta-analysis, with the R package *metafor*, version 1.9-9. A Cohen’s d effect size estimate of the population-level asymmetry was obtained for each hemispheric effect, for each paired left-right measure (Kong et al. 2018). The Cohen’s d hemispheric effects derived from the meta-analytic approach over 99 datasets can be taken as ‘true’ effects representing left-right differences in the average human brain, as measured through this image processing and analysis pipeline. Sixty-three of the hemispheric effects were significant at *p* ≤ 0.05 (uncorrected for multiple testing across regions) in the meta-analysis over 99 datasets, while seven of the effects were not significantly different from zero (*p* > 0.05) (Kong et al. 2018).

### Estimation of Reproducibility

For the present study, each effect within one dataset was compared in turn to the corresponding meta-analytic effect from the 98 other datasets, to avoid sample overlap. For an effect that was significant at *p* ≤ 0.05 in a given meta-analysis of 98 datasets, then the effect in the remaining single dataset would need to be in the same left-right direction as in the meta-analysis, and also be nominally significant within the single dataset (*p* ≤ 0.05), in order to be counted as reproduced in that dataset. For an effect that was non-significant in the meta-analysis (*p* > 0.05), then the effect would also need to be non-significant (*p* > 0.05) in the single dataset, in order to be counted as reproduced in that dataset. As each hemispheric effect was considered as a separate question in this context, no correction for multiple comparisons was applied (uncorrected *p* = 0.05 was used). This ‘yes/no’ dichotomous approach is consistent with the approach that would typically be taken if each dataset had been analyzed in a separate study, to decide whether a given bilateral brain structural measure shows asymmetry or not, and in which left-right direction. Reproducibility was then quantified per effect (i.e., research question) as the proportion of datasets showing hemispheric effects consistent with the meta-analytic effects.

### Reproducibility, Effect Size, and Sample Size

The true hemispheric effect size varied across different brain structural measures (Kong et al. 2018), from Cohen’s *d*=0.0015 to 1.76 (median 0.30) (unsigned magnitudes) (Figure 1). In addition, the sample size varied across datasets, from 14 to 2326 (median 72) (Fig. 1A; Table S1). These variabilities allowed us to examine how reproducibility changes with the true effect size and sample size. As surface area asymmetries are generally more substantial than cortical thickness asymmetries (Fig. 1B) (Kong et al. 2018), we first compared reproducibility rates between the hemispheric effects for these two types of measure. We then calculated the Spearman correlation between the true effect size and the reproducibility rate across all 70 hemispheric effects, as well as across cortical thickness and surface area hemispheric effects separately. We then divided the 70 ‘research questions’ into different groups based on the true effect size: 0.0 ≤ *d* < 0.2, 0.2 ≤ *d* < 0.4, 0.4 ≤ *d* < 0.6, 0.6 ≤ *d* < 0.8, 0.8 ≤ *d<1.8*. Next, we calculated the reproducibility rate for each group, also using a range of minimum sample size thresholds, starting from 15 as in the main analysis above (97 datasets), and then 50 (63 datasets), 100 (37 datasets), 150 (25 datasets), 200 (20 datasets), 300 (19 datasets), and 500 (7 datasets) (Table S1).

**Fig. 1.**
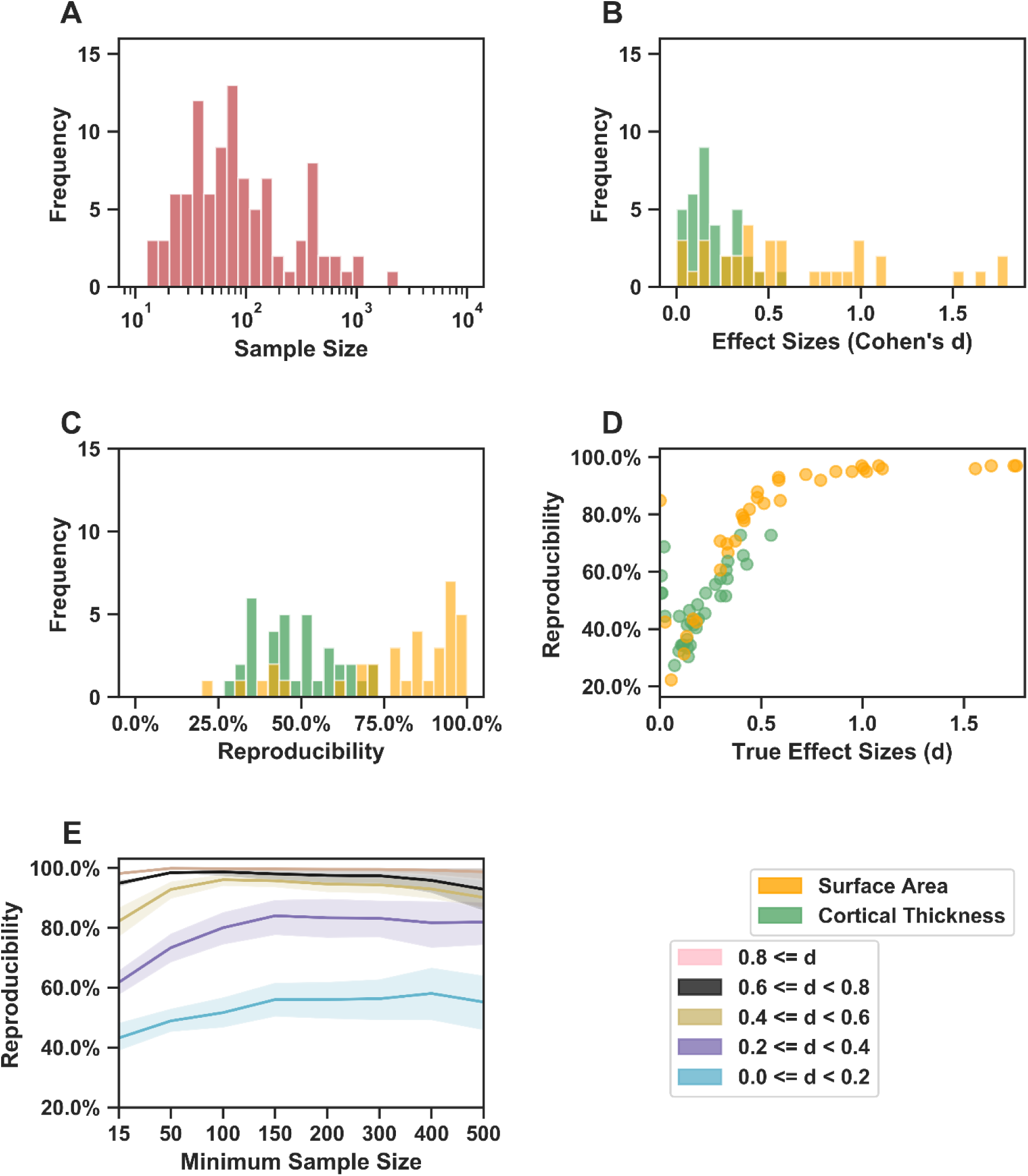
Reproducibility in the absence of selective reporting estimated based on outputs of the ENIGMA cortical asymmetry project. (A). Sample size distribution of the 99 datasets. (B). Effect size distribution of the 70 hemispheric effects of interest. (C). Reproducibility distribution of the 70 hemispheric effects. The reproducibility was assessed by comparing each dataset in turn to the meta-analytic effect from the 98 others, to avoid overlap (see Methods). (D). Scatter plot of the correlation between the reproducibility and the true effect size. (E). Reproducibility changes with both the true effect size and the minimum dataset sample size. Each line plots the mean and 95% confidence interval for reproducibility. We used the meta-analytic effect size over all 99 datasets for visualization purposes. The figure key shows the types of cortical measure (orange indicates surface area; green indicates cortical thickness), as well as groupings by true effect sizes.

### Data and Code Sharing

Data used in this study were from the ENIGMA cortical asymmetry project (http://conxz.net/neurohemi/). Codes for all analyses of the reports are openly shared in GitHub (https://github.com/Conxz/illusReproducibility).

## Results

### Estimating Reproducibility

There was an overall mean reproducibility rate of 63.2%, i.e. 63.2% of tests for hemispheric effects within the separate datasets produced *p* values less than 0.05, together with directions of effect consistent with the true (meta-analytic) effects, or else produced non-significant effects in cases where the true effect was also non-significant. A large variability of reproducibility was observed across effects (*SD* = 22.9%, range from 22.2% to 97.0%) (Fig. 1C). The reproducibility rates were 36.4% and 66.7% for hemispheric effects on average cortical thickness and total surface area respectively (i.e. when assessed for the entire-hemisphere thickness and area measures). For regionally specific hemispheric effects, the reproducibility rate ranged from 27.3% to 72.7% (*Mean* = 48.7%, *SD* = 12.3%), and from 22.2% to 97.0% (*Mean* = 78.4%, *SD* = 21.7%) for cortical thickness and surface area measures, respectively. These findings show that reproducibility is far from perfect, even without any publication bias or potentially problematic practices such as *p*-hacking.

### Reproducibility, Effect Size and Sample size

As expected, regionally specific surface area measures (Fig. 1B) showed significantly higher reproducibility rates for hemispheric effects than cortical thickness measures (Area vs. Thickness: *t(33)* = 6.84, *p* = 3.11e-09) (Fig. 1C), as hemispheric effects on surface area are generally larger (Fig. 1C). Moreover, we found that reproducibility showed a significant correlation with the true effect size for both types of measure (all measures together, *rho* = 0.84, *p* = 2.15e-19; thickness, *rho* = 0.52, *p* = 0.0017; area, *rho* = 0.94, *p* = 4.28e-16) (Fig. 1D). Note that, although this general relation applied to most effects, some true effects very close to zero could also show relatively high reproducibility (Fig. 1D), since in most individual datasets they were found to be low and non-significant as in the meta-analysis (e.g. surface area asymmetry in superior parietal cortex, *d* = 0.002, reproducibility rate = 84.8%, and cortical thickness asymmetry in the pars opercularis, *d* = 0.02, reproducibility rate = 68.7%).

When examining subgroups of effects and datasets according to thresholds on effect size and sample size, we found that the reproducibility rate increased with the minimum sample size threshold, for each specific range of effect size (Fig. 1E). For example, for effects of *d* ≥ 0.6, the reproducibility rate was higher than 90% even when including the datasets with sample sizes as low as 15, while for effects of 0.4 ≤ *d* < 0.6, a minimum sample size of 50 was needed to obtain a reproducibility rate of 90%. Moreover, for effects of 0.2 ≤ *d* < 0.4, a minimum sample size threshold of 100 started to make a reproducibility rate of 80% achievable. In addition, the empirical findings showed that it was impossible to obtain 70% reproducibility for small effects of *d* < 0.2, even with a relatively large minimum sample size threshold of 500.

## Discussion

In this study, we re-visited the summary statistics from a worldwide collaborative neuroscience project (Kong et al. 2018), to empirically assess the reproducibility of realistic biological effects in the absence of *p* hacking or publishing bias, as assessed in heterogeneous neuroscience data and typically-used sample sizes. Overall, reproducibility was limited even in this idealized regime, with a mean rate across all effects = 63.2%, lowest reproducibility rate = 22.2%. Our findings will be useful for guiding future study designs, with respect to anticipated effect sizes and sample sizes.

As expected, the reproducibility rate increased with the true effect size, as well as the sample size of datasets, which together contribute to statistical power. Clearly, to avoid poor reproducibility, a relatively larger sample size is necessary than was available within many of the individual datasets of this study. For example, to obtain a reproducibility rate of 80% for a true effect size of around *d* = 0.4, the sample sizes of individual datasets needed be larger than 100, i.e. greater than the median sample size in this study. There is therefore substantial room to improve reproducibility by increasing sample sizes, even when using currently available methods. Note that the analysis of brain asymmetry involves an inherently paired sample design (i.e. paired left and right measures within subjects), but that the overall picture and principles illustrated here are broadly applicable.

Button et al., (2013) showed that the average statistical power of studies in the neurosciences is low (i.e., around 21%), which is expected to cause low reproducibility, both through false positive and false negative findings. For example, many fMRI studies have traditionally been performed using 10-20 participants (Desmond & Glover 2002). Our observation that reproducibility is strongly influenced by sample size is in line with the PPV calculation (Button et al. 2013) mentioned in the Introduction. However, here we have demonstrated this empirically in a real-world setting, such that *a priori* knowledge of the statistical power (*1*−*β*) and the pre-study odds (*R*) was not necessary. In addition, we have illustrated how reproducibility changes with the true effect size, interacting with sample size. For example, on the basis of our results, if the expected effect size (i.e., Cohen’s *d*) in a paired-measure MRI study is below 0.2, then studies with 500 subjects are still not expected to achieve a reproducibility rate of 80%. Consistent with this, a recent study performed an empirical examination of the replicability of “structural brain behavior” associations using a permutation-based approach (again without any of problems of selective reporting), and concluded that it is relatively unlikely to find an association between behavioral traits and brain morphology with a sample size of less than 500 (replication effect sizes were up to 0.4 (Pearson’s r)) (Kharabian Masouleh et al. 2019).

A reproducibility rate of 36% was reported by the Open Science Framework for 100 findings from psychological studies (Aarts et al. 2015), and a reproducibility rate of 54% for 28 classic findings in psychological science was reported by a more recent Many Lab project (Klein et al. 2018b). Such poor reproducibility has been partly attributed to reporting bias and potentially problematic practices such as selective reporting of outcomes (Aarts et al. 2015; Baker 2016; Bakker et al. 2012; Ioannidis et al. 2014; Ioannidis 2005; Ioannidis 2008; John et al. 2012; Simmons et al. 2011). While we do not dispute the likely relevance of these factors, it is interesting to note that the mean reproducibility rate in the present study, where no such factors were in play, was only 63.2%. As the true effect sizes in the present study ranged from zero to large (Cohen’s *d* up to 1.8), in this respect they can be taken as broadly comparable to those in the human neuroscience and psychology literature, although the effect size distribution within this range might not be representative of the literature at large.

Varying demographic and/or clinical composition of datasets is another factor likely to influence the reproducibility of findings in human neuroscience, even for such fundamental processes as age-related change in neural structure (LeWinn et al. 2017). However, one study that investigated variation in replicability suggested that the contribution of sample heterogeneity can also be modest (Klein et al. 2018b). As noted above and in the Methods, the datasets of the current study differed widely in their age ranges and distributions, and the recruitment criteria were variable too, especially whether subjects were selected as healthy controls without certain psychiatric diseases, or recruited just in the context of unselected population studies. Scanners and scanning parameters differed too (see Table S1). Given this heterogeneity, some ‘non-replication’ could be quite appropriate in certain datasets that comprise specific sub-groups or methodological variants, in which particular effects might be of less relevance. Although this heterogeneity may have contributed to the 63.2% average reproducibility rate, we regard it as a strength of the current study, as we wished the meta-analytic effect sizes to be valid in the context of the heterogeneity typical of the field. Future studies may examine how systematic aspects of MRI dataset heterogeneity influence reproducibility, to gain further insights into the problem that the neuroscience community is facing. In the present study, scanner field strength was somewhat confounded with dataset numbers and sizes, such that we did not investigate this particular aspect of heterogeneity in relation to reproducibility. It is important to note that, while conceptually related to heterogeneity, the reproducibility rate is also influenced by sample and effect sizes, and as such provides a useful method of examining the replicability of effects under real world conditions.

Neuroimaging studies can involve considerable flexibility regarding data processing and statistical analysis, while inconsistent strategies can also contribute to poor reproducibility and contrasting conclusions (Botvinik-Nezer et al. 2019; Pauli et al. 2016). For the present study, the pipeline for MRI quality control, processing and analysis was harmonized, so that the impact of this aspect was necessarily limited. Thus our reproducibility rates may be somewhat idealized, considering how the field typically operates, i.e. with different researchers asking similar questions, but in different datasets and using different strategies. In other words, the reproducibility would likely be worse when the processing pipelines and analysis strategies are different.

Another important aspect affecting reproducibility in the literature may be failure to use blinded designs in primary studies, for example so that researchers know the case-control status of participants while processing their data, and inadvertently introduce bias. This is less likely to be relevant for studies based on automated processing of human brain MRI data, unless there would be bias during visual quality control. As regards our study specifically, the visual quality control of Freesurfer segmentations and parcellations was not done with respect to eventual asymmetry measures, and we have no reason to imagine that the inspection of left and right-hemisphere images was approached differently, on average.

The lowest reproducibility was 22.2%, for a small but significant meta-analytic effect of *d* = 0.052 (cortical area asymmetry in the lingual gyrus). As discussed above, such low reproducibility is likely due to limited power in many of the datasets, to detect such small effects. Below this, there were also 7 nearly-zero, non-significant meta-analytic effects, which were considered to have been reproduced when a given dataset also showed no significant effect. These 7 effects showed relatively higher reproducibility, from 42.4% to 84.8%, than the smallest among the significant meta-analytic effects. In general, this observation highlights the importance of reporting negative, non-significant findings in publications. In addition, as these 7 reproducibility rates were still far from perfect, then dataset heterogeneity and/or technical difficulties in measuring such asymmetries may have been involved.

## Conclusion

Reproducing results is critical for accumulating knowledge in the scientific community. In this study, we re-visited the outputs of a global collaborative neuroscience project (Kong et al. 2018), to empirically demonstrate reproducibility in a real-world setting as regards dataset heterogeneity and sample sizes, but in the absence of *p*-hacking or reporting bias. The results indicate that there is substantial room to improve reproducibility using current neuroimaging methods, even in the absence of *p*-hacking or reporting bias. This can be achieved primarily through increasing statistical power, either through increasing the sample sizes of individual datasets, or via collaborations between researchers, for example in consortia such as ENIGMA (Thompson et al. 2014b). Our study therefore contributes to the ongoing discussion of reproducibility in neuroscience, and science more generally.

## Acknowledgements

Funding information for each site is available in the SI Appendix.

## Conflict of Interest

Any potential conflicts of interest are listed in the SI Appendix.

## Contributions

X.Z.K. conceived this study and performed data analyses; X.Z.K, E.L.W.G, and C.F. contributed data; X.Z.K. and C.F. interpreted the results and wrote the paper. All authors provided feedback on the paper.

