## Supplementary material for "An illustration of reproducibility in neuroscience research in the absence of selective reporting": Table S1

**Abbreviated title:** Reproducibility in the absence of selective reporting


**Authors from the ENIGMA Laterality Working Group**

Xiang-Zhen Kong (1), Samuel R. Mathias (2), Tulio Guadalupe (1), Christoph Abé (3), Ingrid Agartz (4,5,6), Theophilus N. Akudjedu (7), Andre Aleman (8), Saud Alhusaini (9,10), Nicholas B. Allen (11,12), David Ames (13,14), Ole A. Andreassen (15), Alejandro Arias Vasquez (16,17,18,19), Nicola J. Armstrong (20), Phil Asherson (), Felipe Bergo (21), Mark E. Bastin (22,23), Albert Batalla (24), Jochen Bauer (25), Bernhard T Baune (26,27,28), Ramona Baur-Streubel (29), Joseph Biederman (30,31), Sara K. Blaine (2), Premika Boedhoe (32,33,34), Erlend Bøen (6), Anushree Bose (35), Janita Bralten (16,36), Daniel Brandeis (37,38,39,40), Silvia Brem (37,38), Henry Brodaty (41,42), Dilara Yüksel (43), Samantha J. Brooks (44), Jan Buitelaar (36,45,46), Christian Bürger (47), Robin Bülow (48), Vince Calhoun (49,50,51), Anna Calvo (52,53), Erick Jorge Canales-Rodríguez (54,55,56,57), Dara M. Cannon (7), Elisabeth C. Caparelli (58), Francisco X. Castellanos (59,60), Fernando Cendes (21), Tiffany Moukbel Chaim-Avancini (61,62), Kaylita Chantiluke (63), Qun-lin Chen (64,65), Xiayu Chen (66), Yuqi Cheng (67), Anastasia Christakou (68,69), Vincent P. Clark (49,70), David Coghill (71,72), Colm G. Connolly (73,74), Annette Conzelmann (75), Aldo Córdova-Palomera (76), Janna Cousijn (77), Tim Crow (78), Ana Cubillo (63), Udo Dannlowski (47), Sara Ambrosino de Bruttopilo (79), Patrick de Zeeuw (79), Ian J. Deary (80), Damion V. Demeter (81), Adriana Di Martino (59), Erin W Dickie (82), Bruno Dietsche (43), Nhat Trung Doan (76), Colin P. Doherty (83), Alysa Doyle (84,85), Sarah Durston (79), Eric Earl (81), Stefan Ehrlich (86), Carl Johan Ekman (3), Torbjørn Elvsåshagen (87,88), Jeffery N. Epstein (89,90), Damien A. Fair (91,92,93), Stephen V. Faraone (94,95), Guillén Fernández (96,18), Geraldo Busatto Filho (61,62), Katharina Förster (47,97), Jean-Paul Fouche (98), John J. Foxe (99), Thomas Frodl (100), Paola Fuentes-Claramonte (54,55), Janice M. Fullerton (101,102), Hugh Garavan (103), Danielle do Santos Garcia (104), Ian H. Gotlib (105), Anna E. Goudriaan (106,107), Hans Jörgen Grabe (108,109), Nynke A. Groenewold (110), Dominik Grotegerd (47), Oliver Gruber (111), Tiril Gurholt (4), Jan Haavik (94,112), Tim Hahn (47), Narelle K. Hansell (113), Mathew A. Harris (22,114), Catharina A. Hartman (115), Maria del Carmen Valdés Hernández (22,23), Dirk Heslenfeld (116,117), Robert Hester (118), Derrek Paul Hibar (119), Beng-Choon Ho (120), Tiffany C. Ho (74,121), Pieter J. Hoekstra (122), Ruth J. van Holst (123,124), Martine Hoogman (16,36), Marie F. Høvik (125,125), Fleur M. Howells (126), Kenneth Hugdahl (112,127), Chaim Huyser (128,129), Martin Ingvar (3), Akari Ishikawa (104), Anthony James (130), Neda Jahanshad (119), Terry L. Jernigan (131,132), Erik G Jönsson (133,5), Claas Kähler (47,134), Vasily Kaleda (135), Clare Kelly (136,137,138,139), Michael Kerich (140), Matcheri S. Keshavan (141), Sabin Khadka (142), Tilo Kircher (43), Gregor Kohls (143), Kerstin Konrad (143), Ozlem Korucuoglu (144), Bernd Krämer (111), Axel Krug (43), Jonna Kuntsi (145), Jun Soo Kwon (146,147), Nanda Lambregts-Rommelse (17,45), Mikael Landén (148,5), Luisa Lázaro (149,150,151,152), Irina Lebedeva (135), Rhoshel Lenroot (153,154,155), Klaus-Peter Lesch (156,157,158), Qinqin Li (159), Kelvin O. Lim (160), Jia Liu (159), Christine Lochner (161), Edythe D. London (162), Valentina Lorenzetti (163), Michelle Luciano (80), Maartje Luijten (164), Astri J. Lundervold (94,127), Scott Mackey (103), Frank P. MacMaster (165,166,167,168,169), Sophie Maingault (170), Charles B. Malpas (171), Ulrik F. Malt (172,173), David Mataix-Cols (5), Rocio Martin-Santos (174), Andrew R. Mayer (49), Hazel McCarthy (175), Sarah Medland (176), Mitul Metha (177), Philip B. Mitchell (153,178,179), Bryon A. Mueller (160), Susana Muñoz Maniega (22,23), Bernard Mazoyer (180), Colm McDonald (7), Quinn McLellan (181), Katie L. McMahon (182), Genevieve McPhilemy (7), Reza Momenan (140), Angelica M. Morales (162), Janardhanan C. Narayanaswamy (35), José Carlos Vasques Moreira (104), Stener Nerland (6), Liam Nestor (183), Erik Newman (132), Joel T. Nigg (184), Jan Egil Nordvik (185), Stephanie Novotny (142), Eileen Oberwelland Weiss (143), Ruth L. O'Gorman (186,39), Jaap Oosterlaan (187,188), Bob Oranje (79), Catherine Orr (189), Bronwyn Overs (101), Yannis Paloyelis (177), Paul Pauli (29), Martin Paulus (190,191), Kerstin Jessica Plessen (94,192), Georg G. von Polier (143,193), Edith Pomarol-Clotet (54,55), Maria J. Portella (194), Jiang Qiu (64,65), Joaquim Radua (54,55,195,196), Josep Antoni Ramos-Quiroga (197,198), Y.C. Janardhan Reddy (35), Andreas Reif (199), Gloria Roberts (153,178), Pedro Rosa (61,62), Katya Rubia (63), Matthew D. Sacchet (200), Perminder S. Sachdev (41,201), Raymond Salvador (54,55), Lianne Schmaal (11,202,32), Martin Schulte-Rüther (143,203), Lizanne Schweren (122), Jochen Seitz (204), Mauricio Henriques Serpa (61,62), Philip Shaw (205,206), Elena Shumskaya (16,36), Timothy J. Silk (171,207,208), Alan N. Simmons (209,210), Egle Simulionyte (111), Rajita Sinha (2), Zsuzsika Sjoerds (211,212), Runar Elle Smelror (4), Joan Carlos Soliva (197), Nadia Solowij (213), Fabio Luisde Souza-Duran (61,62), Scott R. Sponheim (214), Dan J. Stein (126,215), Elliot A. Stein (58), Michael Stevens (216,217,218), Lachlan T. Strike (113), Gustavo Sudre (205), Jing Sui (49,219), Leanne Tamm (220), Hendrik S. Temmingh (126), Robert J. Thoma (221,222), Alexander Tomyshev (135), Giulia Tronchin (7), Jessica Turner (223), Anne Uhlmann (161,126,103), Theo G.M. van Erp (224), Odile A. van den Heuvel (32,33,34), Dennis van der Meer (225,226), Liza van Eijk (227,113), Alasdair Vance (228), Ilya M. Veer (229), Dick J. Veltman (32), Ganesan Venkatasubramanian (35), Oscar Vilarroya (197,230), Yolanda Vives-Gilabert (231), Aristotle N Voineskos (82,232), Henry Völzke (233,234,235), Daniella Vuletic (126), Susanne Walitza (37,38,39), Henrik Walter (229), Esther Walton (236), Joanna M. Wardlaw (237,238,239), Wei Wen (41), Lars T. Westlye (225,240), Christopher D. Whelan (9), Tonya White (241,242), Reinout W. Wiers (77), Margaret J. Wright (113,243), Katharina Wittfeld (109,108), Tony T. Yang (74), Clarissa L. Yasuda (104), Yuliya Yoncheva (59), Murat Yücel (244), Je-Yeon Yun (245,246), Marcus Vinicius Zanetti (61,247), Zonglei Zhen (66), Xing-xing Zhu (64,65), Georg C. Ziegler (156), Greig I. de Zubicaray (182), Marcel Zwiers (36), Karolinska Schizophrenia Project (KaSP) (248), David C. Glahn (2,249), Fabrice Crivello (180), Simon E. Fisher (1,250), Paul M. Thompson (119), Clyde Francks (1,250)

1. Language and Genetics Department, Max Planck Institute for Psycholinguistics, Nijmegen, The Netherlands.
2. Department of Psychiatry, Yale University School of Medicine, New Haven, CT, USA.
3. Department of Clinical Neuroscience, Osher Centre, Karolinska Institutet, Stockholm, Sweden.
4. Norwegian Centre for Mental Disorders Research (NORMENT), K. G. Jebsen Centre for Psychosis Research, Institute of Clinical Medicine, University of Oslo, Oslo, Norway.
5. Department of Clinical Neuroscience, Centre for Psychiatry Research, Karolinska Institutet, Stockholm, Sweden.
6. Department of Psychiatric Research, Diakonhjemmet Hospital, Oslo, Norway.
7. The Centre for Neuroimaging & Cognitive Genomics (NICOG), Clinical Neuroimaging Lab, NCBES Galway Neuroscience Centre, College of Medicine, Nursing, and Health Sciences, National University of Ireland Galway, H91 TK33 Galway Ireland, Republic of Ireland.
8. BCN Neuroimaging Center, Department of Neuroscience, University Medical Center Groningen, University of Groningen, The Netherlands.
9. Department of Molecular and Cellular Therapeutics, Royal College of Surgeons in Ireland, Dublin, Ireland.

10. Department of Neurology, Yale School of Medicine, Yale University, New Haven, Connecticut, USA.
11. Orygen, The National Centre of Excellence in Youth Mental Health, Parkville, Australia.
12. Department of Psychology, University of Oregon, Eugene OR, USA.
13. National Ageing Research Institute, Melbourne, Australia.
14. Academic Unit for Psychiatry of Old Age, University of Melbourne, Melbourne, Australia.
15. Norwegian Centre for Mental Disorders Research (NORMENT), KG Jebsen Centre for Psychosis Research, Division of Mental Health and Addiction, Oslo University Hospital & Institute of Clinical Medicine, University of Oslo, Oslo, Norway.
16. Department of Human Genetics, Radboud University Medical Center, Nijmegen, The Netherlands.
17. Department of Psychiatry, Radboud University Medical Center, Nijmegen, The Netherlands.
18. Department of Cognitive Neuroscience, Radboud University Medical Center, Nijmegen, The Netherlands.
19. Donders Institute for Brain, Cognition and Behaviour, Radboud University, Nijmegen, The Netherlands.
20. Mathematics and Statistics, Murdoch University, Perth, Australia.
21. Laboratory of Neuroimaging, Department of Neurology, University of Campinas.
22. Centre for Clinical Brain Sciences, University of Edinburgh, Edinburgh, UK.
23. Brain Research Imaging Centre, University of Edinburgh, Edinburgh, UK.
24. Department of Psychiatry, UMC Utrecht Brain Center, University Medical Centre Utrecht, Utrecht University, The Netherlands.
25. Department of Clinical Radiology, School of Medicine, University of Münster, Germany.
26. Department of Psychiatry and Psychotherapy, University of Münster, Germany.
27. Department of Psychiatry, Melbourne Medical School, The University of Melbourne.
28. The Florey Institute of Neuroscience and Mental Health, The University of Melbourne.
29. Department of Psychology, University of Würzburg, Germany, Würzburg, Germany.
30. Department of Psychiatry, Harvard Medical School, Boston, Mass, USA.
31. Clinical and Research Programs in Pediatric Psychopharmacology and Adult ADHD, Massachusetts General Hospital, Boston, MA, USA.
32. Department of Psychiatry, VU University Medical Center, Amsterdam, The Netherlands.
33. Department of Anatomy & Neurosciences, VU University Medical Center, Amsterdam, The Netherlands.
34. Amsterdam Neuroscience, Amsterdam, The Netherlands.
35. Department of Psychiatry, National Institute of Mental Health and Neurosciences, Bengaluru, India.
36. Donders Institute for Brain, Cognition and Behaviour, Nijmegen, The Netherlands.
37. Department of Child and Adolescent Psychiatry and Psychotherapy, University Hospital of Psychiatry Zurich, University of Zurich, Zurich, Switzerland.
38. Neuroscience Center Zurich, University of Zurich and ETH Zurich, Switzerland.
39. Zurich Center for Integrative Human Physiology, University of Zurich, Zurich, Switzerland.
40. Department of Child and Adolescent Psychiatry and Psychotherapy, Central Institute of Mental Health, Medical Faculty Mannheim/Heidelberg University, J5, 68159 Mannheim, Germany.
41. Centre for Healthy Brain Ageing, School of Psychiatry, UNSW Australia.
42. Dementia Collaborative Research Centre ØC Assessment and Better Care, University of New South Wales, Sydney, Australia.
43. Department of Psychiatry and Psychotherapy, Philipps-University Marburg, Germany.
44. School of Natural Sciences and Psychology, Faculty of Science, Liverpool John Moores University, UK.
45. Karakter Child and Adolescent Psychiatry, Nijmegen, The Netherlands.
46. Department of Cognitive Neuroscience, Radboud university, Nijmegen, The Netherlands.
47. Department of Psychiatry, University of Münster, Germany.
48. Department of Diagnostic Radiology and Neuroradiology, University Medicine Greifswald, Greifswald, Germany.
49. The Mind Research Network, Albuquerque, NM, USA.
50. Department of Electrical and Computer Engineering, University of New Mexico, Albuquerque, NM 87131, United States.
51. Tri-institutional Center for Translational Research in Neuroimaging and Data Science (TReNDS) {Georgia State, Georgia Tech, Emory}, Atlanta, GA 30303.
52. Medical Image Core Facility, August Pi I Sunyer Biomedical Research Institute (IDIBAPS), Barcelona, Spain.
53. CIBERBBN.
54. FIDMAG Germanes Hospitalaries Research Foundation, Barcelona, Spain.
55. CIBERSAM, Centro de Investigación Biomédica en Red de Salud Mental, Spain.
56. Department of Radiology, Centre Hospitalier Universitaire Vaudois (CHUV), Lausanne, Switzerland.
57. Signal Processing Laboratory 5 (LTS5), École Polytechnique Fédérale de Lausanne (EPFL), Lausanne, Switzerland.
58. Neuroimaging Research Branch, National Institute on Drug Abuse, National Institutes of Health, Baltimore, Maryland, USA.
59. Department of Child and Adolescent Psychiatry, Hassenfeld Children's Hospital at NYU Langone, New York, USA.
60. Division of Clinical Research, Nathan Kline Institute for Psychiatric Research, Orangeburg, NY, USA.
61. Department of Psychiatry, Faculty of Medicine, University of São Paulo, São Paulo, Brazil.
62. Center for Interdisciplinary Research on Applied Neurosciences (NAPNA), University of São Paulo, São Paulo, Brazil.
63. King's College London, Institute of Psychiatry, Psychology and Neuroscience, Department of Child and Adolescent Psychiatry, London, UK.
64. School of Psychology, Southwest University, Chongqing, China.
65. Key Laboratory of Cognition and Personality, Ministry of Education, Chongqing, China.
66. State Key Laboratory of Cognitive Neuroscience and Learning & IDG/McGovern Institute for Brain Research, Faculty of Psychology, Beijing Normal University, Beijing, China.
67. Department of Psychiatry, First Affiliated Hospital of Kunming Medical University, Kunming, China.
68. Department of Child and Adolescent Psychiatry, Institute of Psychiatry, King's College London, London WC2R 2LS, UK.
69. School of Psychology and Clinical Language Sciences, University of Reading, Reading RG6 6AL, UK.
70. Department of Psychology, University of New Mexico, Albuquerque, NM 87131, United States.
71. Departments of Paediatrics and Psychiatry, University of Melbourne. Victoria, Australia.
72. Division of Neuroscience, Ninewells Hospital and Medical School, University of Dundee.
73. Department of Biomedical Sciences, Florida State University, Tallahassee, FL 32306, USA.
74. Department of Psychiatry, Division of Child and Adolescent Psychiatry, and Weill Institute for Neurosciences, University of California, San Francisco, 401 Parnassus Avenue, San Francisco, CA, USA.
75. Department of Child and Adolescent Psychiatry, Psychosomatics and Psychotherapy, University of Tübingen, Tübingen, Germany.
76. Norwegian Centre for Mental Disorder Research (NORMENT), K.G. Jebsen Centre for Psychosis Research, Division of Mental Health and Addiction, Oslo University Hospital & Institute of Clinical Medicine, University of Oslo, Oslo, Norway.

77. Department of Developmental Psychology, University of Amsterdam, Amsterdam, the Netherlands.
78. SANE POWIC, University Department of Psychiatry, Warneford Hospital, Oxford, UK.
79. NICHE-lab, Brain Center Rudolf Magnus, Department of Psychiatry, University Medical Center Utrecht, Utrecht, The Netherlands.
80. Centre for Cognitive Ageing and Cognitive Epidemiology, Psychology, University of Edinburgh, Edinburgh, UK.
81. Department of Behavioral Neuroscience at Oregon Health & Science University, Portland, OR, USA.
82. Kimel Family Translational Imaging-Genetics Research Laboratory, Campbell Family Mental Health Research Institute, Center for Addiction and Mental Health, Toronto, Canada.
83. Neurology Department, St. James's Hospital, Dublin, Ireland.
84. Department of Psychiatry & Center for Genomic Medicine, Massachusetts General Hospital, Harvard Medical School, Boston, MA,USA.
85. Stanley Center for Psychiatric Research at the Broad Institute, Cambridge, MA, USA.
86. Division of Psychological and Social Medicine and Developmental Neurosciences, Faculty of Medicine, Technische Universität Dresden, Dresden, Germany.
87. Norwegian Centre for Mental Disorder Research (NORMENT), Institute of Clinical Medicine, University of Oslo, Oslo, Norway.
88. Department of Neurology, Oslo University Hospital, Oslo, Norway.
89. University of Cincinnati College of Medicine, Cincinnati, OH, USA.
90. Cincinnati Children's Hospital Medical Center, Cincinnati, OH, USA.
91. Department of Behavioral Neuroscience, Oregon Health & Science University, USA.
92. Department of Psychiatry, Oregon Health & Science University, USA.
93. Advanced Imaging Research Center, Oregon Health & Science University, USA.
94. K.G. Jebsen Centre for Neuropsychiatric Disorders, Department of Biomedicine, University of Bergen, Bergen, Norway.
95. Department of Psychiatry, SUNY Upstate Medical University, Syracuse, NY, USA.
96. Donders Institute for Brain, Cognition and Behaviour, Radboud University Medical Center, Nijmegen, The Netherlands.
97. Institute of Psychiatric Phenomics and Genomics (IPPG), Ludwig-Maximilians-University, Munich, Germany.
98. Department of Psychiatry and Mental Health, University of Cape Town, Cape Town, South Africa.
99. The Ernest J. Del Monte Institute for Neuroscience, Department of Neuroscience, University of Rochester School of Medicine and Dentistry.
100. Department of Psychiatry and Psychotherapy, University Hospital, Otto-von-Guericke-University, Magdeburg, Germany.
101. Neuroscience Research Australia, Sydney, NSW, Australia.
102. School of Medical Sciences, University of New South Wales, Sydney, New South Wales, Australia.
103. Department of Psychiatry, University of Vermont, Burlington, VT, USA.
104. Laboratory of Neuroimaging, Department of Neurology, University of Campinas, Campinas, Brazil.
105. Department of Psychology, Stanford University, USA.
106. Department of Psychiatry, Amsterdam University Medical Center, University of Amsterdam, Amsterdam, The Netherlands.
107. Arkin, Department of Research and Quality of Care & Amsterdam Institute for Addiction Research, Amsterdam, The Netherlands.
108. Department of Psychiatry and Psychotherapy, University Medicine Greifswald, Germany.
109. German Center for Neurodegenerative Diseases (DZNE), Rostock/Greifswald, Germany.
110. UCT Department of Psychiatry & Mental Health, Cape Town, South Africa.
111. Section for Experimental Psychopathology and Neuroimaging, Department of General Psychiatry, Heidelberg University, Heidelberg, Germany.
112. Department of Psychiatry, Haukeland University Hospital, Bergen, Norway.
113. Queensland Brain Institute, University of Queensland, Brisbane, Australia.
114. Division of Psychiatry, University of Edinburgh, Edinburgh, UK.
115. Department of Psychiatry, University Medical Center Groningen, University of Groningen, The Netherlands.
116. Department of Cognitive Psychology, VU University Amsterdam, Amsterdam, The Netherlands.
117. Department of Clinical Neuropsychology, VU University Amsterdam, Amsterdam, The Netherlands.
118. School of Psychological Sciences, University of Melbourne, Melbourne, Australia.
119. Imaging Genetics Center, Mark and Mary Stevens Neuroimaging and Informatics Institute, Keck School of Medicine of USC, Marina del Rey, CA 90292 USA.
120. Department of Psychiatry, University of Iowa College of Medicine, Iowa City, Iowa, USA.
121. Department of Psychology, Stanford University, Stanford, CA, USA.
122. University of Groningen, University Medical Center Groningen, Department of Psychiatry, Groningen, The Netherlands.
123. Amsterdam Institute for Addiction Research, Academic Medical Center, University of Amsterdam, Amsterdam, The Netherlands.
124. Department of Psychiatry, Amsterdam, The Netherlands.
125. Department of Clinical Medicine, University of Bergen, Bergen, Norway.
126. Department of Psychiatry and Mental Health, University of Cape Town, South Africa.
127. Department of Biological and Medical Psychology, University of Bergen, Bergen, Norway.
128. De Bascule, Academic Center for Child and Adolescent Psychiatry, Amsterdam, the Netherlands.
129. AMC, Department of Child and Adolescent Psychiatry, Amsterdam, the Netherlands.
130. University Department of Psychiatry, Warneford Hospital, Oxford, UK .
131. Departments of Cognitive Science, Psychiatry, Radiology, University of California, San Diego, CA, USA.
132. Center for Human Development, University of California, San Diego, CA, USA.
133. Norwegian Centre for Mental Disorders Research (NORMENT), K.G. Jebsen Centre for Psychosis Research, Division of Mental Health and Addiction, Oslo University Hospital, Oslo, Norway.
134. Faculty of Mathematics and Computer Science, University of Münster, Germany.
135. Mental Health Research Center, Moscow, Russia.
136. Department of Psychiatry, University of Dublin, Trinity College Dublin, Dublin, Ireland.
137. The Child Study Center at NYU Langone Medical Center, New York, USA.
138. School of Psychology, Trinity College, Dublin, Ireland.
139. Trinity College Institute of Neuroscience, Dublin, Ireland.
140. Clinical NeuroImaging Research Core, National Institute on Alcohol Abuse and Alcoholism, National Institutes of Health, Baltimore, MD, USA.
141. Beth Israel Deaconess Medical Center, Boston, MA, USA.
142. Olin Neuropsychiatry Research Center, Hartford CT, USA.
143. Child Neuropsychology Section, Department of Child and Adolescent Psychiatry, University Hospital Aachen, Aachen, Germany.
144. Department of Psychiatry, Washington University School of Medicine, St Louis, MO, USA.

145. Social, Genetic and Developmental Psychiatry Centre, Institute of Psychiatry, Psychology and Neuroscience, King's College London, London, UK.
146. Department of Psychiatry, Seoul National University College of Medicine, Seoul, Republic of Korea.
147. Department of Brain and Cognitive Sciences, Seoul National University College of Natural Sciences, Seoul, Republic of Korea.
148. Institute of Neuroscience and Physiology, Sahlgrenska Academy at Gothenburg University, Gothenburg, Sweden.
149. Department of Child and Adolescent Psychiatry and Psychology, Hospital Clínic, Barcelona, Spain.
150. August Pi I Sunyer Biomedical Research Institute (IDIBAPS), Barcelona, Spain.
151. Department of Medicine, University of Barcelona, Barcelona, Spain.
152. CIBERSAM.
153. School of Psychiatry, University of New South Wales, Sydney, NSW, Australia.
154. Neuroscience Research Australia, Sydney, NSW, Australia .
155. University of New Mexico, Albuquerque, New Mexico.
156. Division of Molecular Psychiatry, Center of Mental Health, University of Würzburg, Würzburg, Germany.
157. Laboratory of Psychiatric Neurobiology, Institute of Molecular Medicine, I.M. Sechenov First Moscow State Medical University, Moscow, Russia.
158. Department of Translational Neuroscience, School for Mental Health and Neuroscience (MHeNS), Maastricht University, Maastricht, The Netherlands.
159. Beijing Key Laboratory of Applied Experimental Psychology, National Demonstration Center for Experimental Psychology Education (Beijing Normal University), Faculty of Psychology, Beijing Normal University, Beijing, China.
160. Department of Psychiatry, University of Minnesota, Minneapolis, MN, USA.
161. SU/UCT MRC Unit on Risk and Resilience in Mental Disorders, Department of Psychiatry, Stellenbosch University, South Africa.
162. Department of Psychiatry and Biobehavioral Sciences, University of California, Los Angeles, CA, USA.
163. Institute of Psychology Health and Society, University of Liverpool, Liverpool, UK.
164. Behavioural Science Institute, Radboud University, Nijmegen, The Netherlands.
165. Departments of Psychiatry and Pediatrics, University of Calgary, Calgary AB, Canada.
166. Child and Adolescent Imaging Research Program, Alberta Children's Hospital, Calgary AB, Canada.
167. Mathison Centre for Mental Health Research & Education, Hotchkiss Brain Institute, University of Calgary, Calgary AB, Canada.
168. Strategic Clinical Network for Addictions and Mental Health, Alberta Health Services, Calgary AB, Canada.
169. Alberta Children's Hospital Research Institute, Calgary AB, Canada.
170. Institut des Maladies Neurodégénératives, UMR 5293. Groupe d'Imagerie Neurofonctionnelle, CEA - CNRS - Université de Bordeaux, Bordeaux, France.
171. Developmental Imaging, Murdoch Childrens Research Institute, Royal Children's Hospital, Melbourne, Australia.
172. Department of Research and Education, Oslo University Hospital, Oslo, Norway.
173. Institute of Clinical Medicine, University of Oslo, Oslo, Norway.
174. Department of Psychiatry and Psychology, Hospital Clinic, University of Barcelona, IDIBAPS, CIBERSAM, Barcelona, Spain.
175. Department of Psychiatry, Trinity College Dublin, Dublin, Ireland.
176. Psychiatric Genetics, QIMR Berghofer Medical Research Institute, Brisbane, Queensland, Australia.
177. Department of Neuroimaging, Institute of Psychiatry, Psychology and Neuroscience, King's College London, London, UK.
178. Black Dog Institute, Prince of Wales Hospital, Randwick, NSW, Australia.
179. Prince of Wales Hospital, Sydney, NSW, Australia.
180. Institut des Maladies Neurodégénératives, UMR 5293. Groupe d'Imagerie Neurofonctionnelle, CEA - CNRS - Université de Bordeaux.
181. Faculty of Medicine and Dentistry, University of Alberta, Edmonton, AB, Canada.
182. Institute of Health and Biomedical Innovation, Queensland University of Technology, Brisbane, Australia.
183. Neuropsychopharmacology Unit, Division of Brian Sciences, Imperial College London, London, UK.
184. Department of Psychiatry, Oregon Health & Science University, Portland, OR, USA.
185. Sunnaas Rehabilitation Hospital HT, Nesodden, Norway.
186. Center for MR-Research, University Children's Hospital, Zurich, Switzerland.
187. Emma Children's Hospital, Amsterdam UMC, University of Amsterdam and Vrije Universiteit Amsterdam, Emma Neuroscience Group, department of Pediatrics, Amsterdam Reproduction & Development, Amsterdam, The Netherlands.
188. Vrije Universiteit, Clinical Neuropsychology section, Van der Boechortstraat 7, 1081 BT Amsterdam, Netherlands.
189. Department of Psychological Sciences, Swinburne University of Technology, Melbourne, Australia.
190. Laureate Institute for Brain Research, Tulsa, Oklahoma, USA.
191. Department of Psychiatry, University of California San Diego, La Jolla, California, USA.
192. Child and Adolescent Mental Health Center, Capital Region, Denmark.
193. Institute of Neuroscience and Medicine, Brain & Behaviour (INM-7), Research Centre Jülich, Jülich, Germany.
194. Biomedical Research Institute Sant Pau, Hospital de la Santa Creu i Sant Pau, Centro de Investigación Biomédica en Red de Salut Mental (CIBERSAM), Barcelona, Catalonia, Spain.
195. Centre for Psychiatric Research and Education, Department of Clinical Neuroscience, Karolinska Institutet, Stockholm, Sweden.
196. Department of Psychosis Studies, Institute of Psychiatry, Psychology, and Neuroscience, King's College London, UK.
197. Department of Psychiatry and Legal Medicine, Universitat Autònoma de Barcelona, Barcelona, Spain.
198. Department of Psychiatry, Hospital Universitari Vall d'Hebron, CIBERSAM, Barcelona, Spain.
199. Department of Psychiatry, Psychosomatic Medicine and Psychotherapy, University Hospital Frankfurt, Frankfurt, Germany.
200. Department of Psychiatry and Behavioral Sciences, Stanford University, USA.
201. Neuropsychiatric Institute, Prince of Wales Hospital, Randwick, Australia.
202. Centre for Youth Mental Health, The University of Melbourne, Melbourne, Australia.
203. Translational Brain Research, Department of Child and Adolescent Psychiatry, University Hospital Aachen, Aachen, Germany.
204. Department of Child and Adolescent Psychiatry, University Hospital Aachen, Aachen, Germany.
205. Neurobehavioral Clinical Research Section, National Human Genome Research Institute, Bethesda, USA.
206. National Institute of Mental Health, Bethesda, MD, USA.
207. Department of Paediatrics, University of Melbourne, Melbourne, Australia.
208. School of Psychology, Deakin University, Melbourne, Australia.
209. Department of Psychiatry, University of California, San Diego, 9500 Gilman Dr., La Jolla, CA, USA.
210. Veterans Affairs San Diego Health Care System, La Jolla, CA, USA.
211. Max Planck Institute for Human Cognitive and Brain Sciences, Department of Neurology, Leipzig, Germany.
212. Leiden University, Institute of Psychology, Cognitive Psychology Unit & Leiden Institute for Brain and Cognition, Leiden, The Netherlands.

213. School of Psychology and Illawarra Health and Medical Research Institute, University of Wollongong, Wollongong, Australia.
214. Minneapolis VA Health Care System & University of Minnesota, Minneapolis, MN, USA.
215. SU/UCT MRC Unit on Risk and Resilience in Mental Disorders, Department of Psychiatry and Mental Health, University of Cape Town, South Africa.
216. Clinical Neuroscience and Development Laboratory, Olin Neuropsychiatry Research Center, Hartford CT, USA.
217. Child & Adolescent Research, Hartford Hospital/The Institute of Living, Hartford CT, USA.
218. Department of Psychiatry, Yale University School of Medicine, Hartford CT, USA.
219. National laboratory of Pattern Recognition, Institute of Automation, Chinese Academy of Sciences, Beijing, China.
220. Department of Pediatrics, Division of Behavioral Medicine and Clinical Psychology, Cincinnati Children's Hospital Medical Center, Cincinnati, OH, USA.
221. Department of Psychiatry and Behavioral Sciences, University of New Mexico, Albuquerque, NM, USA.
222. Department of Psychiatry, University of New Mexico, Albuquerque, NM, USA.
223. Department of Psychology and Neuroscience Institute, Georgia State University, Atlanta GA 30302.
224. Department of Psychiatry and Human Behavior, University of California, Irvine, Irvine, USA.
225. NORMENT, KG Jebsen Centre for Psychosis Research, Division of Mental Health and Addiction, Oslo University Hospital & Institute of Clinical Medicine, University of Oslo, Oslo, Norway.
226. School of Mental Health and Neuroscience, Faculty of Health, Medicine and Life Sciences, Maastricht University, The Netherlands.
227. School of Psychology, University of Queensland, Brisbane, Australia.
228. Academic Child Psychiatry Unit, Royal Children's Hospital, University of Melbourne, Melbourne, Victoria, Australia.
229. Charité - Universitätsmedizin Berlin, corporate member of Freie Universität Berlin, Humboldt-Universität zu Berlin, and Berlin Institute of Health, Department of Psychiatry and Psychotherapy, Campus Mitte, Berlin, Germany.
230. Fundació IMIM, Barcelona, Spain.
231. 225 Instituto ITACA, Universitat Politècnica de València, Valencia, Spain.
232. Department of Psychiatry, University of Toronto, Toronto, Canada.
233. Institute for Community Medicine, University Medicine Greifswald, Germany.
234. DZHK (German Centre for Cardiovascular Research), partner site Greifswald, Germany.
235. German Centre for Diabetes Research (DZD), Site Greifswald, Germany.
236. Department of Psychology, Georgia State University, Atlanta GA, USA.
237. Brain Research Imaging Centre, Centre for Clinical Brain Sciences and Dementia Research Institute at the University of Edinburgh, Edinburgh, UK.
238. Scottish Imaging Network, A Platform for Scientific Excellence (SINAPSE) Collaboration, Edinburgh, UK.
239. Centre for Clinical Brain Sciences, Centre for Cognitive Ageing and Cognitive Epidemiology, and UK Dementia Research Institute at The University of Edinburgh, Edinburgh, UK.
240. Department of Psychology, University of Oslo, Oslo, Norway.
241. Department of Child and Adolescent Psychiatry, Erasmus University Medical Centre, Rotterdam, Netherlands.
242. Department of Radiology, Erasmus University Medical Centre, Rotterdam, Netherlands.
243. Centre for Advanced Imaging, University of Queensland, Brisbane, Australia.
244. Monash Institute of Cognitive and Clinical Neurosciences and School of Psychological Sciences, Monash University, Melbourne, Australia.
245. Seoul National University Hospital, Seoul, Republic of Korea.
246. Yeongeon Student Support Center, Seoul National University College of Medicine, Seoul, Republic of Korea.
247. Hospital Sírio-Libanês, São Paulo, Brazil.
248. Members of Karolinska Schizophrenia Project (KaSP) are listed in SI.
249. Olin Neuropsychiatric Research Center, Hartford, CT, USA.
250. Donders Institute for Brain, Cognition and Behavior, Radboud University, Nijmegen, The Netherlands.

**Collaborators from the Karolinska Schizophrenia Project (KaSP) consortium**

Lars Farde (1), Lena Flyckt (1), Göran Engberg (2), Sophie Erhardt (2), Helena Fatouros-Bergman (1), Simon Cervenka (1), Lilly Schwieler (2), Fredrik Piehl (3), Ingrid Agartz (1, 4, 5), Karin Collste (1), Pauliina Victorsson (1), Anna Malmqvist (2), Mikael Hedberg (2), Funda Orhan (2), Carl Sellgren (2, 6)

1. Centre for Psychiatry Research, Department of Clinical Neuroscience, Karolinska Institutet, & Stockholm County Council, Stockholm, Sweden

2. Department of Physiology and Pharmacology, Karolinska Institutet, Stockholm, Sweden

3. Neuroimmunology Unit, Department of Clinical Neuroscience, Karolinska Institutet, Stockholm, Sweden

4. NORMENT, KG Jebsen Centre for Psychosis Research, Division of Mental Health and Addiction, University of Oslo, Oslo, Norway

5. Department of Psychiatry Research, Diakonhjemmet Hospital, Oslo, Norway

6. Centre for Psychiatry Research, Karolinska Institutet, & Stockholm Health Care Services, Stockholm County Council, Karolinska University Hospital, Stockholm, Sweden

**Supporting Information**

- Table S1
- Acknowledgements
- SI Conflicts of interest

### Table S1. Sample information of each dataset.

| DATASETNAME | N | MALE | FEMALE | RIGHT | LEFT | AGEMIN | AGEMAX | AGEMED | ICVMIN | ICVMAX | ICVMED | MALEPROP | RIGHTPROP | FSVERSION | SCANNERFIELD_RAW |
|---|---|---|---|---|---|---|---|---|---|---|---|---|---|---|---|
| **ADHD_ACPU** | 28 | 28 | 0 | NA | NA | 9 | 18 | 13 | 1371960 | 1831540 | 1602485 | 1 | 1 | 5.3 | 3 |
| **ADHD_DUB1** | 41 | 32 | 9 | 32 | 6 | 18 | 49 | 20 | 1114260 | 1958970 | 1705070 | 0.780488 | 0.842105 | 5.3 | 3 |
| **ADHD_WUERZBURG** | 57 | 27 | 30 | 52 | 3 | 24 | 61 | 41 | 1246860 | 1925240 | 1549455 | 0.473684 | 0.945455 | 5.3 | 1.5 |
| **ADHD_RUBIA** | 33 | 33 | 0 | NA | NA | 10 | 18 | 14 | 1304890 | 1886320 | 1570545 | 1 | NA | 5.3 | 3 |
| **ADHD_200KKI** | 65 | 41 | 28 | 61 | 8 | 8 | 13 | 10 | 1194100 | 1731530 | 1483570 | 0.594203 | 0.884058 | 5.3 | 1.5 |
| **ADHD_200NYU** | 111 | 55 | 56 | 107 | 3 | 7 | 18 | 12 | 954388 | 1865180 | 1417110 | 0.495495 | 0.972727 | 5.3 | 3 |
| **ADHD_200OHSU** | 63 | 31 | 39 | NA | NA | 7 | 13 | 10 | 1253320 | 1783890 | 1472580 | 0.442857 | NA | 5.3 | 3 |
| **ADHD_200PEKING** | 143 | 84 | 59 | 141 | 2 | 8 | 15 | 12 | 1162580 | 1862900 | 1494940 | 0.587413 | 0.986014 | 5.3 | 3 |
| **ADHD_AACHEN** | 79 | 53 | 26 | NA | NA | 4 | 17 | 10 | 1176780 | 1771630 | 1486775 | 0.670886 | NA | 5.3 | 3 |
| **ADHD_BERGENADULTS** | 43 | 16 | 27 | 36 | 5 | 21 | 41 | 29 | 1107430 | 1837570 | 1556160 | 0.372093 | 0.878049 | 5.3 | 3 |
| **ADHD_BERGENSVG** | 28 | 20 | 8 | 25 | 2 | 8 | 12 | 10 | 1232950 | 1822780 | 1462155 | 0.714286 | 0.925926 | 5.3 | 3 |
| **ADHD_CAPSUZH** | 36 | 21 | 15 | NA | NA | 8 | 18 | 12.5 | 1137280 | 1835190 | 1576350 | 0.583333 | 0.916667 | NA | 3 |
| **ADHD_DAT_LONDON** | 31 | 31 | 0 | NA | NA | 13 | 19 | 16 | 1169200 | 1506950 | 1340190 | 1 | NA | 5.3 | 3 |
| **ADHD_DUNDEE** | 23 | 10 | 13 | NA | NA | 10 | 18 | 13 | 1190680 | 1773720 | 1461230 | 0.434783 | NA | NA | |
| **ADHD_HARTFORTOLIN** | 109 | 58 | 52 | 106 | 3 | 12 | 19 | 16 | 1018607 | 1736154 | 1375669 | 0.527273 | 0.972477 | NA | |
| **ADHD_IMPACTNL** | 141 | 59 | 82 | 121 | 15 | 19 | 63 | 32 | 1261830 | 1913790 | 1577095 | 0.41844 | 0.889706 | 5.3 | 1.5 |
| **ADHD_MGH** | 69 | 29 | 40 | 63 | 6 | 18 | 59 | 31 | 1116080 | 1811500 | 1451765 | 0.42029 | 0.913043 | 5.1 | 1.5 |
| **ADHD_MTA** | 41 | 31 | 10 | 36 | 3 | 21 | 27 | 24 | 974013 | 1820500 | 1491000 | 0.756098 | 0.923077 | 5.3 | 3 |
| **ADHD_NEUROIMAGEADAM** | 78 | 54 | 24 | 67 | 11 | 12 | 23 | 16 | 1312270 | 1963260 | 1606255 | 0.692308 | 0.858974 | 5.3 | 1.5 |
| **ADHD_NEUROIMAGENIJM** | 39 | 23 | 16 | 34 | 5 | 10 | 23 | 18 | 1298620 | 1965620 | 1632120 | 0.589744 | 0.871795 | 5.3 | 1.5 |
| **ADHD_NICAP** | 81 | 47 | 34 | 69 | 11 | 9 | 11 | 10 | 1326550 | 1877490 | 1595970 | 0.580247 | 0.8625 | 5.3 | 3 |
| **ADHD_NICHE** | 80 | 67 | 13 | 69 | 11 | 7 | 16 | 10 | 1272060 | 1862389 | 1563165 | 0.8375 | 0.8625 | 5.1 | 1.5 |
| **ADHD_NIH** | 304 | 205 | 99 | NA | NA | 4 | 18 | 10 | 1146550 | 1829489 | 1460903 | 0.674342 | NA | 5.1 | 1.5 |
| **ADHD_NYU** | 40 | 22 | 18 | NA | NA | 20 | 49 | 31 | 934852 | 1762720 | 1497460 | 0.55 | 1 | 5.3 | 3 |
| **ADHD_OHSU** | 112 | 60 | 52 | NA | NA | 7 | 13 | 10 | 966984 | 1890000 | 1450000 | 0.535714 | NA | NA | 3 |
| **ADHD_UAB** | 95 | 64 | 31 | NA | NA | 6 | 50 | 27 | 1004270 | 1972030 | 1448310 | 0.673684 | 0.882353 | 5.3 | 3 |
| **ADHD_UCHZ** | 39 | 21 | 18 | NA | NA | 9 | 53 | 15 | 1059950 | 1864990 | 1483240 | 0.538462 | 0.974359 | NA | |

| | | | | | | | | | | | | | | | | |
|---|---|---|---|---|---|---|---|---|---|---|---|---|---|---|---|---|
| **HUBIN_KASP** | 77 | 43 | 34 | 70 | 4 | 20 | 68.97 | 44.85 | 1321762 | 1992782 | 1560192 | 0.558442 | 0.945946 | 5.1,5.3 | 1.5,3 | |
| **TOP1.5T** | 303 | 159 | 144 | 279 | 22 | 18.28 | 73.4 | 33.97 | 1199482 | 2065684 | 1613283 | 0.524752 | 0.92691 | NA | | 1.5 |
| **TOP3T_1** | 380 | 204 | 176 | 277 | 42 | 12.5 | 78 | 31.1 | 1080887 | 2023835 | 1546874 | 0.536842 | 0.868339 | NA | | 3 |
| **TOP3T_2** | 386 | 178 | 208 | 275 | 23 | 12.01 | 88 | 42 | 1190360 | 1981293 | 1545960 | 0.46114 | 0.922819 | NA | | 3 |
| **OCD_LAZARO** | 165 | 92 | 73 | NA | NA | 8 | 17 | 15 | 1211620 | 1999940 | 1549165 | 0.557576 | 0.969697 | 5.3 | 1.5,3 | |
| **CIAM** | 30 | 16 | 14 | 28 | 2 | 19 | 40 | 26.5 | 1070000 | 1790000 | 1425000 | 0.533333 | 0.933333 | 5.3 | | 3 |
| **BIG** | 2326 | 973 | 1353 | 1995 | 103 | 17.39 | 82.66 | 22.47 | 1076480 | 2086070 | 1610760 | 0.418315 | 0.950906 | 5.3 | 1.5,3 | |
| **BIL&GIN** | 453 | 221 | 232 | 248 | 205 | 18.08 | 57.18 | 24.03 | 1111532 | 1835334 | 1396760 | 0.487859 | 0.547461 | 5.3 | | 3 |
| **CAMH** | 146 | 77 | 69 | 139 | 6 | 18 | 86 | 40 | 1123430 | 1833770 | 1471570 | 0.527397 | 0.958621 | 5.3 | | 1.5 |
| **SBP** | 85 | 44 | 41 | NA | NA | 21 | 75 | 35 | 1240000 | 1930000 | 1570000 | 0.517647 | NA | 5.1 | | 1.5 |
| **BRAINN** | 398 | 149 | 249 | 110 | 5 | 18 | 55 | 32 | 811432 | 1900000 | 1330000 | 0.374372 | 0.956522 | 5.3 | | 3 |
| **COLM_UCSF** | 88 | 46 | 42 | NA | NA | 13 | 17 | 15 | 1230000 | 1810000 | 1560000 | 0.522727 | NA | 5.3 | | 3 |
| **CLING** | 323 | 132 | 191 | 307 | 15 | 18 | 58 | 24 | 1062340 | 2081450 | 1581790 | 0.408669 | 0.953416 | 5.3 | | 3 |
| **HMS** | 55 | 21 | 34 | 44 | 7 | 19 | 64 | 41 | 1155270 | 1921180 | 1525650 | 0.381818 | 0.862745 | 5.3 | | 1.5 |
| **COBRE** | 70 | 50 | 20 | NA | NA | 18 | 65 | 34.5 | 1070000 | 1920000 | 1505000 | 0.714286 | 0.985714 | 5.3 | | 3 |
| **MCIC** | 164 | 102 | 62 | 150 | 5 | 18 | 60 | 27 | 1290000 | 1990000 | 1615000 | 0.621951 | 0.967742 | 5.3 | 1.5,3 | |
| **FIDMAG-BARCELONA** | 117 | 55 | 62 | NA | NA | 21 | 63 | 41 | 1211550 | 1850590 | 1509640 | 0.470085 | 1 | 5.3 | | 1.5 |
| **GEB^2** | 674 | 305 | 374 | 616 | 26 | 16.92 | 24.08 | 20.33 | 861850 | 1597510 | 1129565 | 0.44919 | 0.959502 | 5 | | 3 |
| **BIPOLAR KIDS AND SIBS** | 100 | 47 | 53 | 91 | 8 | 14 | 30 | 22 | 995969 | 1784914 | 1360040 | 0.47 | 0.919192 | 5.3 | | 3 |
| **JPCAPETOWN** | 26 | 10 | 16 | NA | NA | 19 | 56 | 29 | 1333950 | 1885170 | 1577910 | 0.384615 | 1 | 5.3 | | 3 |
| **OSLO MALT** | 44 | 18 | 26 | NA | NA | 20 | 50 | 27.5 | 970748 | 1648073 | 1307202 | 0.409091 | 0.977273 | 5.1 | | 3 |
| **STANFORD** | 59 | 23 | 36 | 49 | 9 | 18.85 | 60.53 | 35.73 | 1060000 | 1710000 | 1430000 | 0.389831 | 0.844828 | 5.3 | | 1.5 |
| **CODE** | 74 | 31 | 43 | NA | NA | 20 | 64 | 40.5 | 1300000 | 1930000 | 1570000 | 0.418919 | NA | 5.3 | | 3 |
| **LOTHIAN BIRTH COHORT** | 636 | 336 | 300 | 593 | 38 | 71.04 | 74.22 | 72.69 | 1059294 | 1812082 | 1433641 | 0.528302 | 0.939778 | 5.1 | | 1.5 |
| **FOR2107** | 425 | 160 | 265 | 403 | 22 | 18 | 65 | 27 | 1150870 | 1948960 | 1544530 | 0.376471 | 0.948235 | 5.3 | | 3 |
| **MUENSTER** | 739 | 323 | 416 | 725 | 14 | 17 | 65 | 32 | 930805 | 2015100 | 1432380 | 0.437077 | 0.981055 | 5.3 | | 3 |
| **NESDA** | 65 | 23 | 42 | NA | NA | 21 | 56 | 40 | 992501 | 1906610 | 1499970 | 0.353846 | NA | 5.3 | | 3 |
| **NEUROIMAGE** | 388 | 180 | 208 | NA | NA | 7.72 | 28.59 | 16.53 | 1259250 | 2025730 | 1579145 | 0.463918 | NA | 5.3 | | 1.5 |
| **MAS** | 532 | 242 | 290 | 497 | 16 | 70.3 | 90.06 | 78.08 | 912609 | 1884820 | 1376990 | 0.454887 | 0.968811 | 5.3 | | 3 |
| **OATS** | 413 | 143 | 270 | 369 | 23 | 65 | 89 | 69 | 1052340 | 1882020 | 1425270 | 0.346247 | 0.941327 | 5.3 | 1.5,3 | |

| | | | | | | | | | | | | | | | |
|---|---|---|---|---|---|---|---|---|---|---|---|---|---|---|---|
| **R_SCZ** | 54 | 54 | 0 | NA | NA | 16.07 | 27.62 | 22.94 | 1149026 | 1872418 | 1533725 | 1 | NA | 5.3 | 3 |
| **NUIGALWAY** | 83 | 48 | 35 | 69 | 5 | 18 | 58 | 36 | 1270000 | 1960000 | 1570000 | 0.578313 | 0.932432 | 5.3 | 1.5 |
| **OCD_CHENG_1.5T** | 38 | 11 | 27 | NA | NA | 18 | 49 | 30.5 | 1123950 | 1641540 | 1367960 | 0.289474 | NA | 5.3 | 1.5 |
| **OCD_CHENG_3T** | 93 | 27 | 66 | NA | NA | 22 | 39 | 25 | 933842 | 1753880 | 1307235 | 0.290323 | NA | 5.3 | 3 |
| **OCD_VUMC 1.5T** | 49 | 19 | 30 | 44 | 5 | 21 | 53 | 29 | 1162690 | 1843920 | 1478330 | 0.387755 | 0.897959 | 5.3 | 1.5 |
| **OCD_VUMC 3T** | 36 | 17 | 19 | 29 | 5 | 21.51 | 64.02 | 39.55 | 1271150 | 1780840 | 1567660 | 0.472222 | 0.852941 | 5.3 | 3 |
| **OCD_HUYSER** | 23 | 9 | 14 | 21 | 2 | 8.67 | 16.83 | 14.42 | 1210000 | 1600000 | 1400000 | 0.391304 | 0.913043 | 5.3 | 3 |
| **OCD_MATAIX-COLS** | 33 | 21 | 12 | 17 | 2 | 21 | 63 | 32 | 1200000 | 1970000 | 1450000 | 0.636364 | 0.894737 | 5.3 | 1.5 |
| **OXEOP** | 35 | 16 | 19 | 31 | 4 | 13.7 | 18.88 | 16.27 | 1370000 | 1960000 | 1640000 | 0.457143 | 0.885714 | NA | 1.5 |
| **GBB_ GRADUAL** | 449 | 207 | 242 | NA | NA | 17 | 27 | 20 | 897723 | 1833040 | 1432528 | 0.461024 | NA | 5.3 | 3 |
| **GBB_OLDERS** | 193 | 58 | 135 | NA | NA | 30 | 80 | 60 | 1052606 | 1787736 | 1393392 | 0.300518 | NA | 5.3 | 3 |
| **QTIM** | 1040 | 366 | 674 | NA | NA | 15.4 | 30.11 | 22.04 | 1140000 | 2010000 | 1470000 | 0.351923 | NA | 5.3 | 4 |
| **MACMASTERMDD** | 54 | 23 | 31 | NA | NA | 7 | 24 | 17 | 1280000 | 1880000 | 1570000 | 0.425926 | NA | 5.3 | 1.5 |
| **ESTADO-NARSAD** | 69 | 45 | 24 | NA | NA | 17 | 43 | 27 | 787079 | 1316860 | 1045415 | 0.652174 | NA | 5.3 | |
| **WELLCOME STUDY** | 85 | 45 | 40 | NA | NA | 18 | 50 | 29 | 1014957 | 1860720 | 1465983 | 0.529412 | NA | 5.3 | 1.5 |
| **EPIGEN-IRELAND** | 68 | 39 | 29 | 63 | 5 | 18.35 | 54.11 | 33.44 | 920698 | 2020000 | 1560000 | 0.573529 | 0.926471 | 5.3 | 3 |
| **SHIP** | 448 | 250 | 198 | 408 | 22 | 31 | 89 | 57 | 1215740 | 2036700 | 1573160 | 0.558036 | 0.948837 | 5.3 | 1.5 |
| **SHIP-TREND** | 937 | 528 | 409 | 855 | 36 | 21 | 81 | 51 | 1166230 | 2047120 | 1596805 | 0.563501 | 0.959596 | 5.3 | 1.5 |
| **YOUTH-TOP/NORMENT EOP** | 47 | 20 | 27 | NA | NA | 12.01 | 18.78 | 16.13 | 1288205 | 1880424 | 1531958 | 0.425532 | 0.93617 | 5.3 | 3 |
| **DIP GRONINGEN** | 23 | 6 | 17 | NA | NA | 24 | 66 | 44 | 1207950 | 1816640 | 1471080 | 0.26087 | NA | 5.3 | 3 |
| **NSIOCDS_1.5T_ADULTS** | 20 | 14 | 6 | NA | NA | 18 | 38 | 24.5 | 1084040 | 1685950 | 1424395 | 0.7 | NA | 5.3 | 1.5 |
| **NSIOCDS_3T_ADULTS** | 170 | 108 | 62 | NA | NA | 18 | 45 | 26 | 995985 | 1710000 | 1410000 | 0.635294 | NA | 5.3 | 3 |
| **NSIOCDS_3T_CHILD** | 14 | 7 | 7 | NA | NA | 10 | 18 | 14 | 1140000 | 1670000 | 1340000 | 0.5 | NA | 5.3 | 3 |
| **ADDICTION_COUSIJN** | 40 | 25 | 15 | 37 | 3 | 17.66 | 26.04 | 21.68 | 979900 | 1884780 | 1481670 | 0.625 | 0.925 | 5.3 | 3 |
| **ADDICTION_DSTEIN** | 63 | 50 | 13 | NA | NA | 3 | 53 | 25 | 1170000 | 1860000 | 1600000 | 0.793651 | 1 | 5.3 | 3 |
| **ADDICTION_ESTEIN** | 230 | 113 | 117 | NA | NA | 18 | 55 | 29.5 | 856579 | 1829470 | 1409095 | 0.491304 | 1 | 5.3 | 3 |
| **ADDICTION_FOXE** | 22 | 15 | 4 | NA | NA | 20 | 55 | 40 | 875116 | 1410000 | 1285000 | 0.789474 | NA | 5.3 | 3 |
| **ADDICTION_GARAVAN** | 15 | 11 | 4 | NA | NA | 19 | 33 | 22 | 1230000 | 1700000 | 1470000 | 0.733333 | NA | 5.3 | 3 |
| **ADDICTION_LONDON** | 101 | 45 | 56 | 99 | 2 | 18 | 54 | 33 | 1317580 | 1897910 | 1578800 | 0.445545 | 0.980198 | 5.3 | 1.5 |
| **ADDICTION_LUIJTEN** | 43 | 27 | 16 | NA | NA | 18 | 28 | 22 | 1267940 | 1884160 | 1558980 | 0.627907 | 1 | 5.3 | 3 |

| | | | | | | | | | | | | | | | |
|---|---|---|---|---|---|---|---|---|---|---|---|---|---|---|---|
| **ADDICTION_NIAAA** | 140 | 73 | 67 | 128 | 12 | 21 | 55 | 29 | 880047 | 1870000 | 1380000 | 0.521429 | 0.914286 | 5.3 | 3 |
| **ADDICTION_ORR** | 14 | 13 | 1 | NA | NA | 14 | 19 | 16 | 1459160 | 1816790 | 1654990 | 0.928571 | 1 | 5.3 | 3 |
| **NEURO-ADAPT** | 22 | 12 | 11 | 20 | 1 | 17 | 23 | 20 | 1002660 | 1603610 | 1363665 | 0.521739 | 0.952381 | 5.3 | 3 |
| **ADDICTION_PAULUS** | 31 | 18 | 13 | NA | NA | 18 | 55 | 37 | 1205720 | 1710840 | 1460350 | 0.580645 | NA | 5.3 | 3 |
| **ADDICTION_TRIP** | 18 | 18 | 0 | NA | NA | 29 | 56 | 43 | 1443460 | 1837480 | 1652215 | 1 | 1 | 5.3 | 3 |
| **ADDICTION_SINHA** | 131 | 86 | 45 | 115 | 15 | 19 | 50 | 27 | 844969 | 1350000 | 1090000 | 0.656489 | 0.884615 | 5.3 | 3 |
| **ADDICTION_NESDA-AD** | 20 | 14 | 6 | 17 | 3 | 27 | 68 | 49.5 | 1260000 | 1850000 | 1565000 | 0.7 | 0.85 | 5.3 | 3 |
| **ADPG** | 24 | 24 | 0 | NA | NA | 21 | 53 | 38.5 | 1295890 | 2013820 | 1674265 | 1 | 1 | 5.3 | 3 |
| **ADDICTION_YUCEL** | 179 | 116 | 66 | NA | NA | 18 | 55 | 19 | 1290000 | 1990000 | 1590000 | 0.637363 | NA | 5.3 | 1.5,3 |
| **SEOUL III** | 89 | 54 | 35 | NA | NA | 18 | 48 | 25 | 1190000 | 1920000 | 1570000 | 0.606742 | NA | 5.3 | 3 |
| **SEOUL II** | 103 | 57 | 46 | NA | NA | 18 | 36 | 24 | 1200000 | 1830000 | 1550000 | 0.553398 | NA | 5.3 | 1.5 |
| **SEOUL I** | 45 | 29 | 16 | NA | NA | 18 | 42 | 24 | 1250000 | 1920000 | 1560000 | 0.644444 | NA | 5.3 | 1.5 |

**Acknowledgements**

**ENIGMA Center.** P.M.T., N.J., and D.P.H. were supported in part by a grant from the NIH Big Data to Knowledge (BD2K) Program (U54 EB020403).

**Addiction_Cousijn.** This study investigated the predictive role of neurocognitive functions in the progression from cannabis use to dependence in at-risk young adults. JC & AG received funding for the Cannabis Prospective study from ZonMW grant no.31180002 from the Netherlands Organization for Scientific Research (NWO)

**Addiction_DStein.** The Meth-CT studies investigate structural and functional brain alterations in methamphetamine-dependent individuals compared to healthy controls, and the neural underpinnings of psychotic symptoms. Research was supported by the Department of Psychiatry and Mental Health and the Human Research Ethics Committee, University of Cape Town, and the Medical Research Council, South Africa.

**Addiction_EStein.** Data collection was supported by the Intramural Research Program of NIDA/NIH.

**Addiction_Foxe.** These projects studied cognitive control and reward processing in current and abstinent cocaine users. HG & JF received funds from NIDA: R01-DA014100

**Addiction_Garavan.** This data was supported by USPHS grant from the National Institute on Drug Abuse: DA01865-01, Australian Research Council Grant (RH) DP0556602 and Australian National Health and Medical Research Council Career Development Award 519730 (RH).

**Addiction_London.** These projects examined how structural and functional brain abnormalities were associated with attention, working memory (R01DA015179, EDL), response inhibition, cognitive flexibility and decision making in methamphetamine users (R01DA020726, P20DA022539, EDL). Additional support for these projects came from the Thomas P. and Katherine K. Pike Chair in Addiction Studies and the Endowment from the Marjorie Greene Family Trust (EDL).

**Addiction_Luijten.** ML & DV received funding for the DABIS study from VIDI grant no.016.08.322 from the Netherlands Organization for Scientific Research (NWO) awarded to Ingmar H A Franken.

**Addiction_NESDA-AD.** ZS & DV received funding for the NESDA-AD study from ZonMW grant no. 31160004 from the Netherlands Organization for Scientific Research (NWO).

**Addiction_NIAAA.** Data collection by RM was supported by the Intramural Clinical and Biological Research (DICBR) Program of the National Institute on Alcohol Abuse and Alcoholism (NIAAA), National Institutes of Health.

**Addiction_Orr.** The study was approved by the School of Psychology in Trinity College Dublin and was conducted in accordance with the declaration of Helsinki.

**Addiction_Paulus.** MP received funding from NIMH: R01 DA018307

**Addiction_Sinha.** Rajita Sinha received funds from NIH/NIDA: P50-DA016556, R01-AA013892, UL1-DE019586, PL1-DA024859

**Addiction_TrIp.** LS & DV received funding for the TrIP study from ZonMW grant no. 31160003 from the Netherlands Organization for Scientific Research (NWO).

**Addiction_Yucel.** This study was funded by National Health and Medical Research Council (NHMRC) of Australia (Project Grant 459111 to Nadia Solowij). MY was supported by a National Health and Medical Research Council Fellowship (#1117188) and the David Winston Turner Endowment Fund. NS was supported by an Australian Research Council Future Fellowship (FT110100752). This study has been done in part with Spanish grants: Plan Nacional sobre Drogas,

Ministerio de Sanidad y Consumo PNSD/2011/050 (IP: R. Martin-Santos) and PNSD2006/101 (IP: R. Martin-Santos); and the support of DIUE of Generalitat de Catalunya SGR2009/1435.

**ADPG.** AG & RvH received funding for the ADPG study from ZonMW grant no.91676084 from the Netherlands Organization for Scientific Research (NWO).

**ADHD_Wuerzburg**. This work was supported by the German Research Foundation (DFG; KFO 125/2, project 7 to PP).

**BIG.** The Brain Imaging Genetics (BIG) database was established in Nijmegen in 2007. This resource is now part of Cognomics, a joint initiative by researchers of the Donders Centre for Cognitive Neuroimaging, the Human Genetics and Cognitive Neuroscience departments of the Radboud University Medical Center, and the Max Planck Institute for Psycholinguistics. The Cognomics Initiative is supported by the participating departments and centres and by external grants, i.e. the Biobanking and Biomolecular Resources Research Infrastructure (Netherlands) (BBMRI-NL), the Hersenstichting Nederland, and the Netherlands Organisation for Scientific Research (NWO). The research on BIG also receives funding from the European Community‘s Seventh Framework Programme (FP7/2007–2013) under grant agreements #602450 (IMAGEMEND) and #602805 (Aggressotype) and from the National Institutes of Health (NIH) Consortium grant U54 EB020403, supported by a cross-NIH alliance that funds Big Data to Knowledge Centers of Excellence. We would like to thank all persons who kindly participated in this research. In addition, AF Marquand gratefully acknowledges support from the Language in Interaction project, funded by the NWO under the Gravitation Programme (grant 024.001.006).

**BIL&GIN.** The BIL&GIN was designed to allow an in-depth exploration of hemispheric specialization and of its variability in human. A local ethics committee (CCPRB Basse-Normandie) approved the experimental protocol.

**Bipolar Kids and Sibs.** This study and research team are supported by the Australian National Medical and Health Research Council (programme grant 1037196; project grants 1066177 and 1063960), the Lansdowne Foundation and the Janette Mary O'Neil Research Fellowship (to JMF).

**BRAINN.** The Brazilian Institute of Neuroscience and Neurotechnology (BRAINN) was launched in 2013 by FAPESP (SÃO PAULO RESEARCH FOUNDATION, grant 2013/07559-3) as a Research, Innovation and Dissemination Center (RIDC).

**CAMH.** The CAMH dataset was collected in Toronto with support from the CAMH Foundation and the Canadian Institutes of Health Research.

**CIAM.** The CIAM study was conducted at the University of Cape Town, Department of Psychiatry and Mental Health, and was supported by the Department of Psychiatry and Mental Health and University Research Committee, University of Cape Town, South Africa and the National Research Foundation South Africa.

**CliNG.** Recruitment for the CliNG study sample was partially supported by the Deutsche Forschungsgemeinschaft (DFG) via the Clinical Research Group 241 'Genotype-phenotype relationships and neurobiology of the longitudinal course of psychosis', TP2 (PI Gruber; http://www.kfo241.de; grant number GR 1950/5-1).

**CODE.** The CODE cohort was collected from studies funded by Lundbeck and the German Research Foundation (WA 1539/4-1, SCHN 1205/3-1, SCHR 443/11-1).

**Colm_UCSF.** This work was supported by the Brain and Behavior Research Foundation grant (formerly NARSAD) to T.T.Y. and by a US National Institute of Mental Health (NIMH) grant to T.T.Y. (R01MH085734).

**DIP GRONINGEN.** Data collection for DIP, as contributed to ENIGMA projects, was funded by the Gratama Foundation, the Netherlands.

**EPIGEN-Ireland.** The work was supported by research grants from the Science Foundation Ireland (Research Frontiers Program award 08/RFP/GEN1538) and Brainwave–the Irish Epilepsy Association.

**ESTADO-NARSAD.** The present investigation was supported by a 2010 NARSAD Independent Investigator Award (NARSAD: The Brain and Behavior Research Fund) awarded to Geraldo F. Busatto. Geraldo F. Busatto is also partially funded by CNPq-Brazil. Marcus V. Zanetti is funded by FAPESP, Brazil (no. 2013/03905-4).

**GBB.** This research was supported by the National Natural Science Foundation of China (31271087; 31470981; 31571137; 31500885), National Outstanding young people plan, the Program for the Top Young Talents by Chongqing, the Fundamental Research Funds for the Central Universities (SWU1509383,SWU1509451), Natural Science Foundation of Chongqing (cstc2015jcyjA10106), Fok Ying Tung Education Foundation (151023) , General Financial Grant from the China Postdoctoral Science Foundation (2015M572423, 2015M580767), Special Funds from the Chongqing Postdoctoral Science Foundation (Xm2015037), Key research for Humanities and social sciences of Ministry of Education(14JJD880009).

**GEB^2.** The project was supported by National Natural Science Foundation of China (31230031, 31221003, 31471067, 31470055).

**JPCapeTown.** This work was supported by the Medical Research Council of South Africa, the Obsessive-Compulsive Foundation (Dan J. Stein), the National Research Foundation of South Africa (Christine Lochner), and an unrestricted grant from Lundbeck H/S, and we acknowledge the contribution of our research assistants.

**Lothian Birth Cohort.** Data collection was supported by the Disconnected Mind project, funded by Age UK.  J.M.W. is partly funded by the Scottish Funding Council as part of the SINAPSE Collaboration. The work was undertaken by The University of Edinburgh Centre for Cognitive Ageing and Cognitive Epidemiology, part of the cross-council Lifelong Health and Wellbeing Initiative (MR/K026992/1). Funding from the Biotechnology and Biological Sciences Research Council (BBSRC) and MRC is gratefully acknowledged. We thank the study participants. We also thank Catherine Murray for recruitment of the participants and the radiographers and other staff at the Brain Research Imaging Centre.

**MacMasterMDD.** Funding from the Halifax Stanley Centre. Support for this research in part from the Cuthbertson and Fischer Chair in Paediatric Mental Health, the Alberta Children's Hospital Foundation, Alberta Children's Hospital Research Institute for Child and Maternal Health, the Mathison Centre for Mental Health Research & Education, the Hotchkiss Brain Institute, and the University of Calgary.

**MAS.** We would like to acknowledge and thank the Sydney MAS participants, their supporters and the Sydney MAS Research Team. Sydney MAS is supported by the National Health and Medical Research Council (NHMRC) Program Grants (350833, 56896, 109308).

**MCIC.** This work was supported primarily by the Department of Energy DE-FG02-99ER62764 through its support of the Mind Research Network (MRN, formerly known as the MIND Institute) and the consortium as well as by the National Association for Research in Schizophrenia and Affective Disorders (NARSAD) Young Investigator Award (to SE) as well as through the Blowitz-Ridgeway and Essel Foundations, and through NWO ZonMw TOP 91211021, the DFG research fellowship (to SE), the Mind Research Network, National Institutes of Health through NCRR 5MO1-RR001066 (MGH General Clinical Research Center), NIMH K08 MH068540, the Biomedical Informatics

Research Network with NCRR Supplements to P41 RR14075 (MGH), M01 RR 01066 (MGH), NIBIB R01EB006841 (MRN), R01EB005846 (MRN), 2R01 EB000840 (MRN), 1RC1MH089257 (MRN), as well as grant U24 RR021992, P20RR021938/P20GM103472 and R01MH094524.

**NESDA.** The infrastructure for the NESDA study (http://www.nesda.nl) is funded through the Geestkracht program of the Netherlands Organisation for Health Research and Development (Zon-Mw, grant no 10-000-1002) and is supported by participating universities and mental health care organizations (VU University Medical Center, GGZ inGeest, Arkin, Leiden University Medical Center, GGZ Rivierduinen, University Medical Center Groningen, Lentis, GGZ Friesland, GGZ Drenthe, Scientific Institute for Quality of Healthcare (IQ healthcare), Netherlands Institute for Health Services Research (NIVEL) and Netherlands Institute of Mental Health and Addiction (Trimbos Institute).

**Neuro-ADAPT.** OK received support for the Neuro-ADAPT study from VICI grant no. 453.08.01 from the Netherlands Organization for Scientific Research (NWO) awarded to Reinout W Wiers.

**NeuroIMAGE.** This work was supported by NIH Grant R01MH62873 (to Stephen V. Faraone), NWO Large Investment Grant 1750102007010 and NWO Brain & Cognition an Integrative Approach grant (433-09-242) (to Jan Buitelaar), and grants from Radboud University Nijmegen Medical Center, University Medical Center Groningen and Accare, and VU University Amsterdam. The research leading to these results also received funding from the European Community’s Seventh Framework Programme (FP7/2007– 2013) under grant agreement numbers 278948 (TACTICS), 602450 (IMAGEMEND) and n° 602805 (Aggressotype), and from the European Community’s Horizon 2020 Programme (H2020/2014 – 2020) under grant agreement n° 643051 (MiND). Barbara Franke is supported by a Vici grant from NWO (grant number 016-130-669). In addition, Jan Buitelaar and Barbara Franke are supported by a grant for the ENIGMA Consortium (grant number U54 EB020403) from the BD2K Initiative of a cross-NIH partnership.

**NSIOCDS_1.5T_Adults.** The structural MRI data were obtained as part of three funded projects Government of India grants to the Wellcome-DBT India Alliance grant to Dr. Venkatasubramanian (500236/Z/11/Z), Prof. Reddy (SR/S0/HS/0016/2011) and Dr. Narayanaswamy (DST INSPIRE faculty grant -IFA12-LSBM-26) of the Department of Science and Technology; the Government of India grants to Prof. Reddy No.BT/PR13334/Med/30/259/2009) and Dr. Narayanaswamy (BT/06/IYBA/2012) of the Department of Biotechnology.

**NSIOCDS_3T_Adults.** The structural MRI data were obtained as part of three funded projects Government of India grants to the Wellcome-DBT India Alliance grant to Dr. Venkatasubramanian (500236/Z/11/Z), Prof. Reddy (SR/S0/HS/0016/2011) and Dr. Narayanaswamy (DST INSPIRE faculty grant -IFA12-LSBM-26) of the Department of Science and Technology; the Government of India grants to Prof. Reddy No.BT/PR13334/Med/30/259/2009) and Dr. Narayanaswamy (BT/06/IYBA/2012) of the Department of Biotechnology.

**NSIOCDS_3T_Child.** The structural MRI data were obtained as part of three funded projects Government of India grants to the Wellcome-DBT India Alliance grant to Dr. Venkatasubramanian (500236/Z/11/Z), Prof. Reddy (SR/S0/HS/0016/2011) and Dr. Narayanaswamy (DST INSPIRE faculty grant -IFA12-LSBM-26) of the Department of Science and Technology; the Government of India grants to Prof. Reddy No.BT/PR13334/Med/30/259/2009) and Dr. Narayanaswamy (BT/06/IYBA/2012) of the Department of Biotechnology.

**NUIG.** This NUI Galway study was supported by the NUI Galway Millennium Fund and grant funding from the Health Research Board (HRA_POR/2011/100).

**OATS.** We would like to acknowledge and thank the OATS participants, their supporters and the OATS Research Team. OATS is supported by a National Health and Medical Research Council

(NHMRC)/Australian Research Council Strategic Award (Grant 401162) and the NHMRC Project Grant (1045325). OATS was facilitated through access to the Australian Twin Registry, which is funded by the NHMRC Enabling Grant 310667.

**OCD_Cheng_1.5T.** This study was supported the Funding of Yunnan Provincial Health Science and Technology Plan (2010NS016, 2011WS008), the united founding of Yunnan Administration of Science & Technology and Kunming Medical College(2011FB167).

**OCD_Cheng_3T.** This study was supported by grants from National Natural Science Foundation of China (NSFC) (81101005), the Ministry of Science and Technology of Yunnan Province(2012FB158), the Funding of Yunnan Provincial Health Science and Technology Plan (2014NS171, 2014NS172), the united founding of Yunnan Administration of Science & Technology and Kunming Medical College(2011FB167).

**OCD_Huyser.** The studies were supported by a grant from the Amsterdam school of neuroscience (ONWA) for scan costs.

**OCD_Lazaro.** The studies were supported by two grants from Marato_TV3 Foundation (01/2010, 091710).

**OCD_Mataix-Cols.** These structural scans come from a series of studies conducted at King's College London and funded by the Wellcome Trust (Mary L Phillips, PI) and a pump priming grant from the South London and Maudsley Trust, London (project grant no. 064846; David Mataix-Cols PI).

**OCD_VUmc 1.5T.** Supported by the Dutch Organization for Scientific Research (NWO) (grants 912-02-050, 907-00-012, 940-37-018, and 916.86.038).

**OCD_VUmc 3T.** Supported in part by the Netherlands Society for Scientific Research (NWO-ZonMw VENI grant 916.86.036 to Dr. van den Heuvel; NWO-ZonMw AGIKO stipend 920-03-542 to Dr. de Vries), and a NARSAD Young Investigators Award to Dr. van den Heuvel, Amsterdam Brain Imaging Platform to Dr. van den Heuvel, the Netherlands Brain Foundation (2010(1)-50 to Dr. van den Heuvel).

**Oslo Malt.** The study is funded by the Research Council of Norway (167153/V50, 204966/F20), the South-Eastern Norway Regional Health Authority, Oslo University Hospital, and research grants from Mrs. Aslaug Throne-Holst and from the Ebbe Frøland Foundation.

**OXEOP.** MRC funded grant number: G0500092 - Anatomical connectivity in early onset schizophrenia

**QTIM.** QTIM is funded by the National Institutes of Health (project ROI HD HD050735; NIH Award 1U54EB020403-01, subaward no. 56929223) and the NHMRC (1009064, 496682). Ethics approval was given by the Human Research Ethics Committees of the Queensland Institute of Medical Research, University of Queensland, and Uniting Health Care. We thank the twins and siblings for their participation, Marlene Grace and Ann Eldridge for twin recruitment, Aiman Al Najjar and other radiographers for scanning, and Kerrie McAloney and Daniel Park for research support.

**R_SCZ.** The R_SCZ database has been supported by MHRC and by a research grant from The Russian Foundation for Basic Research (grant code 15-06-05758 A; grantee Dr. Irina Lebedeva, PhD, DrSci (biol), the head of the Laboratory of Neuroimaging and Multimodal analysis, MHRC).

**SBP.** The SBP is supported by grants from the Swedish Medical Research Council (K2014-62X-14647-12-51, K2010-61P-21568-01-4, and K2013-61X-08276-26-4), the Swedish foundation for Strategic Research (KF10-0039), the Swedish Brain foundation (FO2016-0176), and the Swedish Federal Government under the LUA/ALF agreement (ALFGBG-426721).

**Seoul I.** This study was supported by the Korean Research Foundation (1998-003-F00172), Korean Health Research and Development Grant (HMP-98-N-2-0029), Korea Research Foundation Grant (KRF-2001-044-F00182), Korean Research Foundation (2001-041-F00182), Seoul National University Hospital Research Fund (11-2003-001), and Brain Research Center of the 21st Century Frontier Research Program by Ministry of Science and Technology of Republic of Korea (M103KV010007 04K2201 007 10).

**Seoul II.** This study was supported by grants (M103KV010012-06K2201-01210, 2009K001270, and 2010K000817) from Brain Research Center of the 21st Century Frontier Research Program funded by the Ministry of Science and Technology of the Republic of Korea, a grant (M10644020003-08N4402-00310) from the Cognitive Neuroscience Program of the Korean Ministry of Science and Technology of the Republic of Korea, the Korea Research Foundation grants funded by the Korean Government (KRF-2007-313-E00306 and KRF-2008-313-E00341), World Class University program through the Korea Science and Engineering Foundation funded by the Ministry of Education, Science and Technology (R31-10089, and R32-10142), a grant from the Seoul National University Hospital Research Fund (04-2008-104), and a grant from the National Research Foundation of Korea (2012-0005150) funded by the Ministry of Education, Science and Technology (MEST) of the Republic of Korea.

**Seoul III.** This study was supported by National Research Foundation of Korea grant funded by the Ministry of Education, Science and Technology (MEST) of the Republic of Korea (2011-0015639 and 2012-0005150), a grant of the Korea Health Technology R&D Project, Ministry of Health & Welfare of the Republic of Korea (A110094), and Basic Science Research Program through the National Research Foundation of Korea (NRF) funded by the Ministry of Science, ICT and Future Planning (2013R1A2A1A03071089).

**SHIP** and **SHIP-TREND** are part of the Community Medicine Research Network of the University Medicine Greifswald, which is supported by the German Federal State of Mecklenburg- West Pomerania. MR imaging was further supported by Siemens Healthineers, Erlangen, Germany.

**Stanford.** The Stanford dataset was established with the support of NIMH Grant R01MH59259 to Ian Gotlib, and the National Science Foundation Integrative Graduate Education and Research Traineeship (NSF IGERT) Recipient Award 0801700 and National Science Foundation Graduate Research Fellowship Program (NSF GRFP) DGE-1147470 to Matthew Sacchet.

**Wellcome Study.** This study was funded by the Wellcome Trust, UK.

**Youth-TOP/NORMENT EOP.** Funding is provided by the Norwegian Research Council (NFR), the South-Eastern Norway Regional Health Authority and the KG Jebsen Foundation.

**TOP3T_2.** The TOP study was supported by the Research Council of Norway (#160181, 190311, 223273, 213837, 249711), the South-East Norway Health Authority (2014114, 2014097, 2017- 112), and the Kristian Gerhard Jebsen Stiftelsen (SKGJ-MED-008) and the European Community's Seventh Framework Programme (FP7/2007–2013), grant agreement no. 602450 (IMAGEMEND).

**TOP1.5T.** The TOP study was supported by the Research Council of Norway (#160181, 190311, 223273, 213837, 249711), the South-East Norway Health Authority (2014114, 2014097, 2017- 112), and the Kristian Gerhard Jebsen Stiftelsen (SKGJ-MED-008) and the European Community's Seventh Framework Programme (FP7/2007–2013), grant agreement no. 602450 (IMAGEMEND).

**HUBIN_KASP.** KaSP was supported by the Swedish Research Council (K2015-62X-15077-12-3), and by grants from the Swedish Medical Research Council (SE: 2009-7053; 2013-2838; SC: 523-2014-3467), the Swedish Brain Foundation, Åhlén-siftelsen, Svenska Läkaresällskapet, Petrus och Augusta Hedlunds Stiftelse, Torsten Söderbergs Stiftelse, the AstraZeneca-Karolinska Institutet Joint Research Program in Translational Science, Söderbergs Königska Stiftelse, Professor Bror Gadelius

Minne, Knut och Alice Wallenbergs stiftelse, Stockholm County Council (ALF and PPG), Centre for Psychiatry Research, KID-funding from the Karolinska Institutet. The HUBIN study was supported by the Swedish Research Council (grant numbers K2015-62X-15077-12-3), the regional agreement between Karolinska Institutet and Stockholm County Council, the Karolinska Institutet and the Knut and Alice Wallenberg Foundation.

**Muenster.** This work was funded by the German Research Foundation (SFB-TRR58, Project C09 to UD) and the Interdisciplinary Center for Clinical Research (IZKF) of the medical faculty of Münster (grant Dan3/012/17 to UD).

**FOR2107.** This work was funded by the German Research Foundation (DFG, grant FOR2107 KI 588/14-1 to TK, KO4291/3-1 to AK and DA1151/5-1 to UD).

**FIDMAG-Barcelona.** This work was supported by the Catalonian Government (2014-SGR-1573) and by the Plan Nacional de I+D+i 2008–2011 and 2013–2016: Juan de la Cierva-formación contract (FJCI-2015-25278 to PF-C). Also by the Instituto de Salud Carlos III and co-funded by European Union (ERDF/ESF, "Investing in your future"): Miguel Servet Research Contracts (MS14/00041 to JR and CPII16/00264 to EP-C) and Research Project Grants (PI15/00277 to EC-R, PI11/01766 and PI14/00292 to JR, PI14/01148 to EP-C and PI14/01151 to RS).

**ADHD-ACPU.** Scans taken as part of National competitive research grant funding awarded to Alasdair Vance and Timothy Silk.

**ADHD_NICAP.** The study was funded by the National Medical Health and Research Council of Australia (NHMRC; project grant #1065895).

**ADHD_OHSU.** The OHSU dataset was established through several Foundation grants and grants from the National Institutes of Health: R01 MH115357 (MPI: Fair, Nigg), R56 MH086654 (MPI: Nigg, Fair), R01 MH086654 (PI: Nigg), R01 MH099064 (PI: Nigg), R01 MH096773 (PI: Fair), DeStefano Family Innovation Fund (PI: Fair), R00 MH091238 (PI: Fair), Oregon Clinical and Translational Research Institute (UL1TR000128).

**ADHD_UCHZ.** This work was supported by the University Research Priority Program "Integrative Human Physiology" at the University of Zurich.

**ADHD_Dundee.** This work was supported by a Tenovus-Scotland initiative (a local trust) and by SINAPSE (www.sinapse.ac.uk), which included a SINAPSE-SPIRIT industry partnership with Siemens Medical (a SINAPSE studentship for Blair Johnston).

**TOP3T_1.** The TOP study was supported by the Research Council of Norway (#160181, 190311, 223273, 213837, 249711), the South-East Norway Health Authority (2014114, 2014097, 2017- 112), and the Kristian Gerhard Jebsen Stiftelsen (SKGJ-MED-008) and the European Community's Seventh Framework Programme (FP7/2007–2013), grant agreement no. 602450 (IMAGEMEND).

**ADHD_IMPACTNL.** This study was supported by grants from the Netherlands Organization for Scientific Research (NWO), i.e. the NWO Brain & Cognition Excellence Program (grant 433-09-229) and a Vici grant to BF (grant 016-130-669), a Veni grant (no. 91619115) to MH, and by grants from the Netherlands Brain Foundation (grant 15F07[2]27) and BBMRI-NL (grant CP2010-33). The research leading to these results also received funding from the European Community's Seventh Framework Programme (FP7/2007 – 2013) under grant agreements n° 602805 (Aggressotype) and n° 602450 (IMAGEMEND), and from the European Community's Horizon 2020 Programme (H2020/2014 – 2020) under grant agreement n° 643051 (MiND). In addition, the work was supported by a grant for the

ENIGMA Consortium (grant number U54 EB020403) from the BD2K Initiative of a cross-NIH partnership.

**ADHD_MTA.** Data collection was funded in part by the National Institute on Drug Abuse (Contract #: HHSN271200800009C).

**COBRE.** This research was supported by NIH1R01-EB006841, NIH1R01-EB005846, NIH2R01-EB000840, NIH1 P20 RR021938-01 and DOEDEFG02-08ER64581 (to VDC); the national high tech development plan (863 plan) 2015AA020513 (to JS); R01 MH65304 and VA CSR&D IIR-04-212-3 (to JMC). TW is supported by the Netherlands Organization for Health Research and Development (ZonMw) TOP project number 91211021 and the Simons Foundation Autism Research Initiative (SFARI - 307280).

**SI Conflicts of interest**

The ENIGMA co-authors declare no conflicts of interest except for the authors below:

*Theo Van Erp* consulted for Roche Pharmaceuticals and has a contract with Otsuka Pharmaceutical, Ltd.

*Anders Dale* is a Founder of CorTechs Labs, Inc. He serves on the Scientific Advisory Boards of CorTechs Labs and Human Longevity, Inc., and receives research funding through a Research Agreement with General Electric Healthcare.

Stephen Faraone received income, potential income, travel expenses continuing education support and/or research support from Lundbeck, KenPharm, Rhodes, Arbor, Ironshore, Shire, Akili Interactive Labs, CogCubed, Alcobra, VAYA, Sunovion, Genomind and NeuroLifeSciences. With his institution, he has US patent US20130217707 A1 for the use of sodium-hydrogen exchange inhibitors in the treatment of ADHD.

*Paulo Mattos* was on the speakers' bureau and/or acted as consultant for Janssen-Cilag, Novartis, and Shire in the previous five years; he also received travel awards to participate in scientific meetings from those companies. The ADHD outpatient program (Grupo de Estudos do Déficit de Atenção/Institute of Psychiatry) chaired by Dr. Mattos has also received research support from Novartis and Shire.The funding sources had no role in the design and conduct of the study; collection, management, analysis, or interpretation of the data; or preparation, review, or approval of the manuscript.

*Tobias Banaschewski* served in an advisory or consultancy role for Hexal Pharma, Lilly, Medice, Novartis, Oxford outcomes, PCM scientific, Shire and Viforpharma. He received conference support or speaker's fee by Janssen McNeil, Lilly, Medice, Novartis and Shire. He is/has been involved in clinical trials conducted by Shire & Viforpharma. The present work is unrelated to the above grants and relationships.

*Katya Rubia* received speaker's fees form Shire, Medice and a grant from Lilly for another project.

*Jan Haavik* has received speaker fees from Lilly, Novartis and Janssen Cilag.

*Steve Faraone* has received income, travel expenses and/or research support from, and/or has been on an Advisory Board for, and/or participated in continuing medical education programs sponsored by:

Pfizer, Ironshore, Shire, Akili Interactive Labs, CogCubed, Alcobra, VAYA Pharma, Neurovance, Impax, NeuroLifeSciences, Otsuka, McNeil, Janssen, Novartis, Eli Lilly and the NIH. With his institution, he has US patent US20130217707 A1 for the use of sodium-hydrogen exchange inhibitors in the treatment of ADHD. He receives royalties from books published by Guilford Press: Straight Talk about Your Child's Mental Health; Oxford University Press: Schizophrenia: The Facts; Elsevier, ADHD: Non-Pharmacologic Treatments

*Kerstin Konrad* received speaking fees from Medice, Lilly and Shire.

*Josep-Antoni Ramos* was on the speakers' bureau and/or acted as consultant for Eli-Lilly, Janssen-Cilag, Novartis, Shire, Lundbeck, Almirall and Rubió in the last 3 years. He also received travel awards (air tickets + hotel) for taking part in psychiatric meetings from Janssen-Cilag, Rubió, Shire, and Eli- Lilly. The ADHD Program chaired by him received unrestricted educational and research support from the following pharmaceutical companies in the last 3 years: Eli-Lilly, Rovi, Ferrer, Lundbeck, Shire, and Rubió.

*Pieter Hoekstra* received a research grant from Shire and was part of the advisory board of Shire.

*Jan Buitelaar* has been in the past 3 years a consultant to / member of advisory board of / and/or speaker for Janssen Cilag BV, Eli Lilly, Medice, Shire, Roche, and Servier. He is not an employee of any of these companies, and not a stock shareholder of any of these companies. He has no other financial or material support, including expert testimony, patents, royalties.

*David Coghill* has been in the past 3 years a consultant to / member of advisory board of / and/or speaker for Janssen Cilag, Eli Lilly, Medice, Shire, Novartis. He receives royalties from Oxford University Press. He is not an employee of any of these companies, and not a stock shareholder of any of these companies.

*D.P.H.* is now a Senior Scientist for Janssen, Inc., but his work for this manuscript was completed while he was a faculty member at USC.

*Dr. Joseph Biederman* is currently receiving research support from the following sources: AACAP, The Department of Defense, Food & Drug Administration, Headspace, Lundbeck, Neurocentria Inc., NIDA, PamLab, Pfizer, Shire Pharmaceuticals Inc., Sunovion, and NIH.

*Dr. Biederman* has a financial interest in Avekshan LLC, a company that develops treatments for attention deficit hyperactivity disorder (ADHD). His interests were reviewed and are managed by Massachusetts General Hospital and Partners HealthCare in accordance with their conflict of interest policies.

*Dr. Biederman's program* has received departmental royalties from a copyrighted rating scale used for ADHD diagnoses, paid by Ingenix, Prophase, Shire, Bracket Global, Sunovion, and Theravance; these royalties were paid to the Department of Psychiatry at MGH.

*In 2017, Dr. Biederman* is a consultant for Aevi Genomics, Akili, Guidepoint, Ironshore, Medgenics, and Piper Jaffray. He is on the scientific advisory board for Alcobra and Shire. He received honoraria from the MGH Psychiatry Academy for tuition-funded CME courses. Through MGH corporate licensing, he has a US Patent (#14/027,676) for a non-stimulant treatment for ADHD, and a patent pending (#61/233,686) on a method to prevent stimulant abuse.

*In 2016, Dr. Biederman* received honoraria from the MGH Psychiatry Academy for tuition-funded CME courses, and from Alcobra and APSARD. He was on the scientific advisory board for Arbor Pharmaceuticals. He was a consultant for Akili and Medgenics. He received research support from Merck and SPRITES.

*In 2015, Dr. Biederman* received honoraria from the MGH Psychiatry Academy for tuition-funded CME courses, and from Avekshan. He received research support from Ironshore, Magceutics Inc., and Vaya Pharma/Enzymotec.

*In 2014, Dr. Biederman* received honoraria from the MGH Psychiatry Academy for tuition-funded CME courses. He received research support from AACAP, Alcobra, Forest Research Institute, and Shire Pharmaceuticals Inc.

*In previous years, Dr. Biederman* received research support, consultation fees, or speaker's fees for/from the following additional sources: Abbott, Alza, APSARD, AstraZeneca, Boston University, Bristol Myers Squibb, Cambridge University Press, Celltech, Cephalon, The Children's Hospital of Southwest Florida/Lee Memorial Health System, Cipher Pharmaceuticals Inc., Eli Lilly and Co., Esai, ElMindA, Fundacion Areces (Spain), Forest, Fundación Dr.Manuel Camelo A.C., Glaxo, Gliatech, Hastings Center, Janssen, Juste Pharmaceutical Spain, McNeil, Medice Pharmaceuticals (Germany), Merck, MGH Psychiatry Academy, MMC Pediatric, NARSAD, NIDA, New River, NICHD, NIMH, Novartis, Noven, Neurosearch, Organon, Otsuka, Pfizer, Pharmacia, Phase V Communications, Physicians Academy, The Prechter Foundation, Quantia Communications, Reed Exhibitions, Shionogi Pharma Inc, Shire, the Spanish Child Psychiatry Association, The Stanley Foundation, UCB Pharma Inc., Veritas, and Wyeth.

*Henry Brodaty* is on the Advisory Board for Nutricia and has conducted an Alzheimer's drug trial for Tau Therapeutics.
